# Stepwise firing mechanism of an extracellular contractile injection system

**DOI:** 10.1101/2025.10.08.681156

**Authors:** Jingwei Xu, Charles F. Ericson, Elena R. Toenshoff, Martin Pilhofer

## Abstract

Contractile injection systems (CISs) mediate cell-cell interactions and are widespread among bacteria and archaea. These phage tail-like macromolecular machines puncture their target by a tube that is propelled by a contractile sheath. The mechanism underlying CIS firing, which starts with target binding and ends with sheath contraction, remains unclear.

Here, we investigate the extracellular CIS from *Algoriphagus machipongonensis* (AlgoCIS) by a multimodal cryo-electron microscopy approach and structure-guided engineering, which allowed us to arrest AlgoCIS in multiple intermediate states of firing. Together with the post-firing structure, our data suggest a stepwise firing mechanism involving all modules: signal propagation starts with the binding of the tail-fibers, followed by opening of the cage, an expansion of the baseplate iris, and resulting in sheath contraction and the release of cap adaptor.

Our study will serve as a framework for understanding the firing mechanism of diverse CISs and will facilitate the engineering of CISs for biomedical applications.

## Introduction

Bacteria evolved numerous types of sophisticated macromolecular complexes to mediate interactions with bacterial, archaeal, or eukaryotic cells^1,2^. Contractile injection systems (CISs), translocate effector proteins into the extracellular space or directly into the target organism^3–9^. The overall structure of CISs is conserved and resembles the contractile tail of phage T4, comprising a baseplate complex, a helical array of a contractile sheath, and a conduit inner tube capped by a spike complex^10–12^. The firing of CISs is thought to be triggered through contact with a target cell, resulting in conformational changes of the baseplate and contraction of the sheath, eventually propelling the inner tube across the target envelope^11,13,14^. The mechanism of the coupling of target binding to sheath contraction is poorly understood, in particular because it has so far been impossible to capture any intermediate structures of this dynamic process. For the pyocin R2, a ‘checksum’ mechanism has been proposed, in which the tail-fibers pull on the baseplate wedges upon binding to initiate the firing^15^.

Besides these pyocin R2-like assemblies, genome-wide bioinformatic analyses revealed a phylogenetically distinct group of CISs that are widespread among bacterial and archaeal phyla^16–18^. Interestingly, these closely related systems function by distinct modes of action, including extracellular CIS (eCIS)^3,19–21^, the membrane-bound type VI secretion system (T6SS)^5,6,22^, cytoplasmic CIS (CIS^Sc^)^23,24^, and the thylakoid membrane-anchored CIS (tCIS)^25^. In some cases, individual CIS complexes can also further assemble into larger super-structures, as seen for clusters of the T6SS*^iv^* clusters in endosymbiotic bacteria^22^ or arrays of metamorphosis-associated contractile structures from marine bacteria^3^. Among this group of CISs, eCIS is the most common mode of action, by which the particles are assembled in the bacterial cytoplasm, released upon cell lysis and then bind to their target cell surface^3,19^. This eCIS mode of action also shows tremendous potential for biomedical applications through programmable engineering^26–28^.

Here we set out to investigate the firing mechanism of an eCIS representative from the marine bacterium *Algoriphagus machipongonensis* PR1 (AlgoCIS)^19^. This strain is genetically tractable, and expresses large amounts of AlgoCIS that carries two cargo proteins in the inner tube lumen. Cryo-electron microscopy (cryoEM) analyses discovered multiple novel but conserved structural components, such as a baseplate cage surrounding the central spike, and a cap adaptor that connects the cap with the distal sheath layer (Extended Data Fig. 1). A previous low-resolution structure of the post-firing state revealed an intact baseplate iris ring^19^, suggesting a firing mechanism that may be distinct from pyocin R2. The propagation of the signal through the different eCIS modules that leads to the eventual sheath contraction is unclear.

## Results

### Cap and sheath-tube modules undergo significant structural rearrangements upon firing

We started by characterizing the structure of the post-firing AlgoCIS. We purified AlgoCIS particles, triggered firing via guanidine-HCl (GdCl) treatment, determined the cryoEM structure, and built an atomic model (Fig. 1a, Extended Data Fig. 2, and Extended Data Table 1). To verify this post-firing model, we also determined a cryoEM structure from a sample where AlgoCIS firing was triggered by low pH (Extended Data Fig. 3a-c and Extended Data Table 1), showing a high overall agreement with the structure from the GdCl-treated sample (Extended Data Fig. 3d-e).

**Figure 1.**
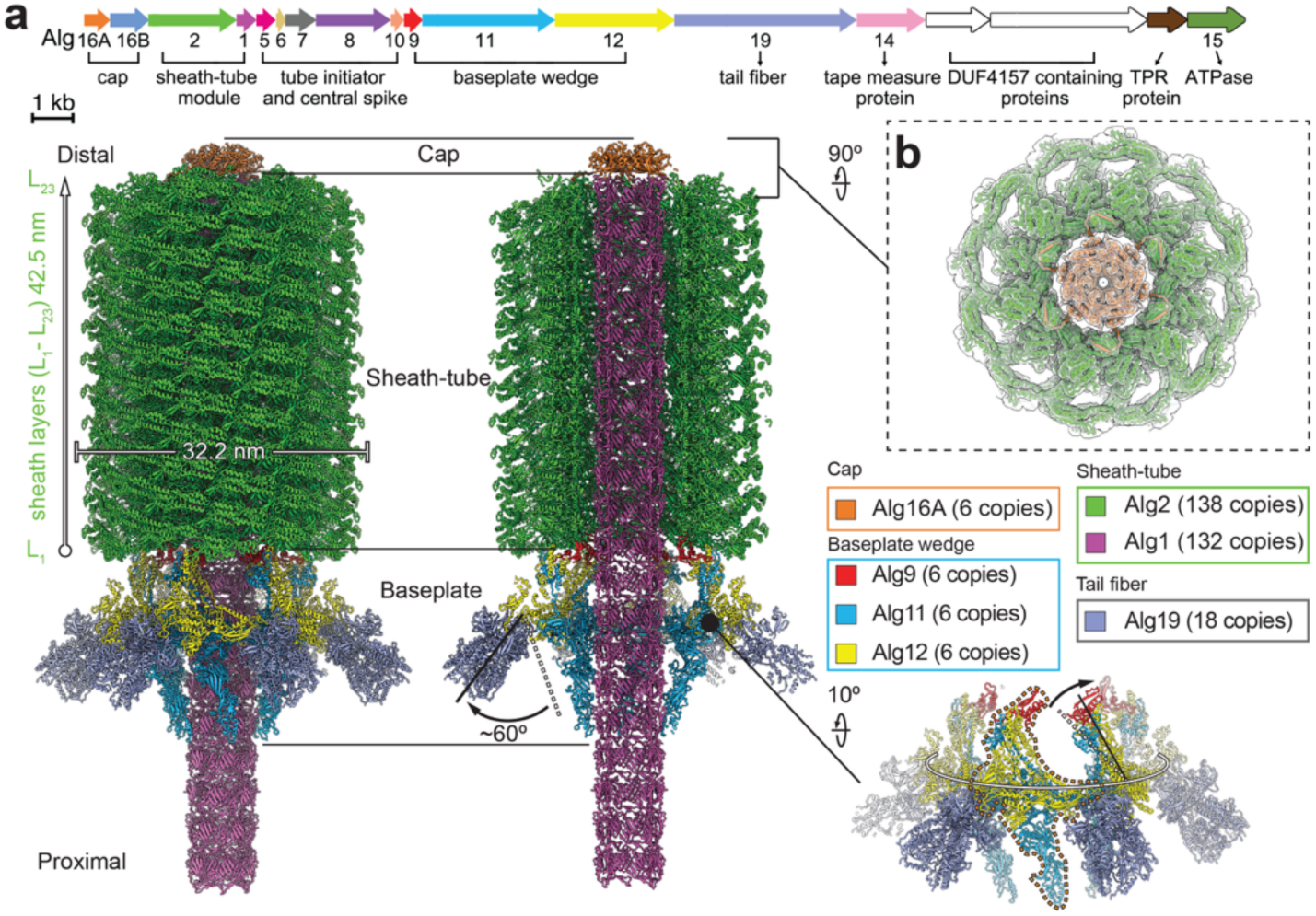
AlgoCIS undergoes significant structural re-organization upon firing. **a.** Atomic model of AlgoCIS in the post-firing state (left: side view; middle: central sliced view), showing structural re-arrangements upon firing. The sheath is contracted, the tube is propelled forward, the baseplate iris is expanded, the cage is opened, and the tail-fibers are tilted outwards. The AlgoCIS gene cluster is shown on top. The structure of the baseplate in the post-firing AlgoCIS is shown on the bottom right and the structural rearrangements are indicated by arrows. One baseplate wedge is outlined by a dashed line. **b.** Top view of the cap module in the post-firing AlgoCIS, showing that only the cap protein is observed and the cap adaptor protein (Alg16B) is not present in the post-firing structure. The EM density is shown transparent.

Like AlgoCIS in the pre-firing state^19^ (Extended Data Fig. 1), the post-firing structure of AlgoCIS comprises three modules: cap, sheath-tube, and baseplate (Fig. 1a). Among them, the sheath-tube module adopts the most pronounced re-arrangement — it contracts from 92.4 nm to 42.5 nm in length, with the outer diameter increasing from 24.6 nm to 32.2 nm (Fig. 1a). As reported for other CISs^6,20,21,23,29–32^, sheath contraction disrupts contacts with the inner tube, which is propelled forward to puncture the target cell membrane.

The distal end of the contracted sheath is terminated by six copies of the cap protein (Alg16A), whose C-terminal loop rotates ∼21° to accommodate the sheath expansion (Extended Data Fig. 4a). As a result, the distance between the topmost sheath layer and the cap protein (Alg16A) decreases upon firing (36 Å *vs.* 70 Å in the pre-firing state) (Extended Data Fig. 4b). Moreover, the shortened distance does not provide enough space for the binding of the cap adaptor Alg16B, which is forced to dissociate upon sheath contraction. Accordingly, we did not observe Alg16B in the post-firing structure (Fig. 1b), and mass spectrometry analyses detected fewer hits for Alg16B in the sample of post-firing AlgoCIS (Extended Data Fig. 4c). Note that in a AlgoCISΔAlg16B knockout mutant several types of aberrant assemblies (e.g. tube-baseplate complexes, contracted with jammed tube particles, or incomplete CISs) have been observed previously^19^. Together, these data indicate that the cap adaptor functions to stabilize and lock AlgoCIS in the pre-firing state.

### Baseplate iris expands and remains intact by tail-fibers upon firing

The proximal end of the contracted sheath connects to the baseplate that is formed by six baseplate wedges (Fig. 1a), each consisting of a sheath initiator Alg9 and a heterodimer of Alg11/12 (Fig. 2a). Six heterodimers of Alg11/12 together assemble into an iris-ring like structure (hereafter called “iris”, Fig. 2b). Structural comparison with the pre-firing baseplate revealed various degrees of outward movements of individual domains of Alg11/12, which together contribute to a slight expansion of the iris (Fig. 2b and Extended Data Fig. 5a-b). Domain II of Alg11, which holds the central spike in the pre-firing state, rotates ∼29° to enlarge the inner diameter of the iris to provide space for the translocation of the tube (Fig. 2b and Extended Data Fig. 5b). The Alg11 domains III and IV perform a similar rotation to open the cage upon firing (Extended Data Fig. 5b). Alg9 docks onto the core bundle of the heterodimer Alg11/12 and follows the outward tilt of Alg11/12 upon firing (Fig. 2a), resulting in an expansion (26 Å) and rotation (∼35°) of the Alg9 ring (Fig. 2c and Extended Data Fig. 5c). Given the extensive contacts between Alg9 and the first sheath layer (Extended Data Fig. 5d), the structural rearrangements of Alg9 introduce a twist force onto the sheath, thereby likely initiating its contraction as shown in other CISs and phages^13,15,33^.

**Figure 2.**
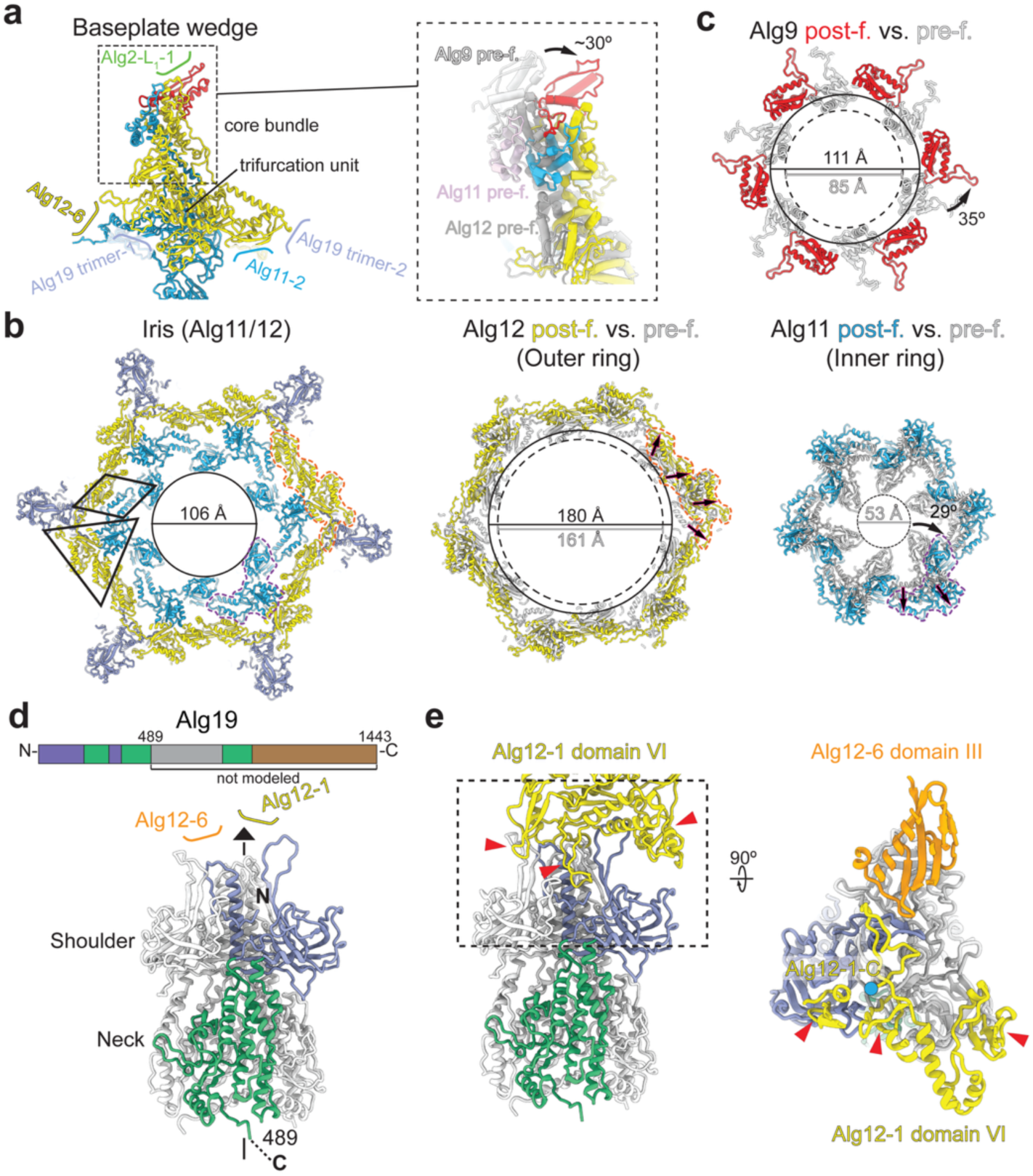
Baseplate iris expands and remains intact by tail-fibers. **a.** Each baseplate wedge undergoes an outward tilt upon firing. Core bundle and trifurcation unit on the baseplate wedge are labeled. Zoom-in of the core bundle is shown on the right, which tilts ∼30° outwards compared to the one in the pre-firing state (Alg9: white; Alg11: pink; Alg12: grey). **b.** The iris remains intact in the post-firing AlgoCIS. Shown are the overall structure of the baseplate iris ring in the post-firing state (left) and individual comparisons of the outer ring (middle) and inner ring (right) before (white) and after firing. Single baseplate proteins are outlined with colored dashed lines (Alg11: purple; Alg12: red). The trifurcation unit and lateral dimer are highlighted by a triangle or a box, respectively. The outward movements and rotations of individual domains in one baseplate wedge are indicated by arrows. **c.** Alg9 ring expands and rotates (indicated by an arrow) upon firing. Shown is a structural comparison of Alg9 in the pre- (white) and post-firing (red) state. The rotation of an individual protein is indicated. **d.** Three copies of Alg19 form one tail-fiber in AlgoCIS. Top: Alg19 domain organization. Bottom: the cryoEM structure of one tail-fiber including shoulder and neck domains; one protomer is color-coded. The central three-fold axis is represented by a triangle. **e.** One tail-fiber connects two adjacent baseplate wedges (see **b,** left panel). The left panel shows that three protrusion loops (red arrowheads) from domain VI of Alg12 (Alg12-1: yellow) bind to the shoulder domains of one tail-fiber. The right panel shows the top view of the boxed area. Note that the C-terminal triangular loop in Alg12 (Alg12-1) binds to three protomers of the tail-fiber, while domain III from the adjacent Alg12 (Alg12-6: orange) interacts with the top part of one protomer. The C-terminus of Alg12-1 is represented by blue circle.

Interestingly, the iris structure remains intact in the post-firing state (Fig. 2b), with the six vertices being bound to one tail-fiber each. Each tail-fiber consists of a homotrimer of Alg19 and Alphafold^34^ predicts four structural domains: shoulder, neck, middle and Ig-rich domain, respectively (Extended Data Fig. 6a). Like other structural modules in AlgoCIS, tail-fibers perform substantial conformational re-arrangements upon firing –– the shoulder and neck domains of the tail-fiber undergo an outward tilt (∼60°) with a pivot point on the binding site with the iris (Fig. 1a).

To investigate the interaction between the iris and the tail-fiber, we determined the structure of the N-terminal part of the tail-fiber (including shoulder and neck domains) via focused refinement (Fig. 2d, Extended Data Fig. 2a-c). The Alg19 shoulder folds similar to some phage receptor binding proteins (e.g. lactococcal phage 1358 and p2) and also similar to TssK from T6SSs (Extended Data Fig. 6b-c). The contact between tail-fiber and iris is mainly mediated by three central α-helices of the Alg19 shoulder domain that interact with the C-terminal loop of Alg12 (Fig. 2e and Extended Data Fig. 6d). This is reminiscent of the complex TssG-TssK seen in T6SSs^35^. The tail-fiber attachment is further stabilized via additional contacts: three protrusion loops of one Alg12 hold two protomers of the tail-fiber, and the adjacent Alg12 binds to the third protomer (Fig. 2e). The tail-fibers therefore bind to the vertices of the iris by bridging two neighboring baseplate wedges. Notably, this binding profile is similar to that seen between the iris and the crown in tCIS^25^ (Extended Data Fig. 6e), which also shows an intact iris upon firing. On the other hand, this is in stark contrast to Afp^21,36^ and pyocin R2 (ref.^15^), where each tail-fiber interacts with only one baseplate wedge and the iris dissociates upon firing (Extended Data Fig. 6e).

### Tail-fibers mediate AlgoCIS binding to bacterial cells

Next, we explored the structure and possible roles of the tail-fibers in the AlgoCIS firing mechanism. As some eCIS have been reported to bind to targets via their tail-fibers^15,26,28^, we started out imaging purified AlgoCIS in the presence of bacterial cells by cryo-electron tomography (cryoET). We co-incubated AlgoCIS with the marine bacterium *Echinicola pacifica*, a strain that was previously co-isolated with *A. machipongonesis* PR1 (ref.^37^). We observed that some AlgoCIS particles were loosely (non-perpendicularly) attached to the bacterial surface via the flexible C-terminal part of the tail-fibers (Extended Data Fig. 7a). Notably, we also detected many surface-bound, perpendicularly oriented particles in the pre-firing (867 particles in 112 tomograms) and in the post-firing (47 particles in 112 tomograms, Fig. 3a and Extended Data Fig. 7b) states. The latter represents the actual puncturing event, where the inner tube penetrates the bacterial outer membrane (Fig. 3a).

**Figure 3.**
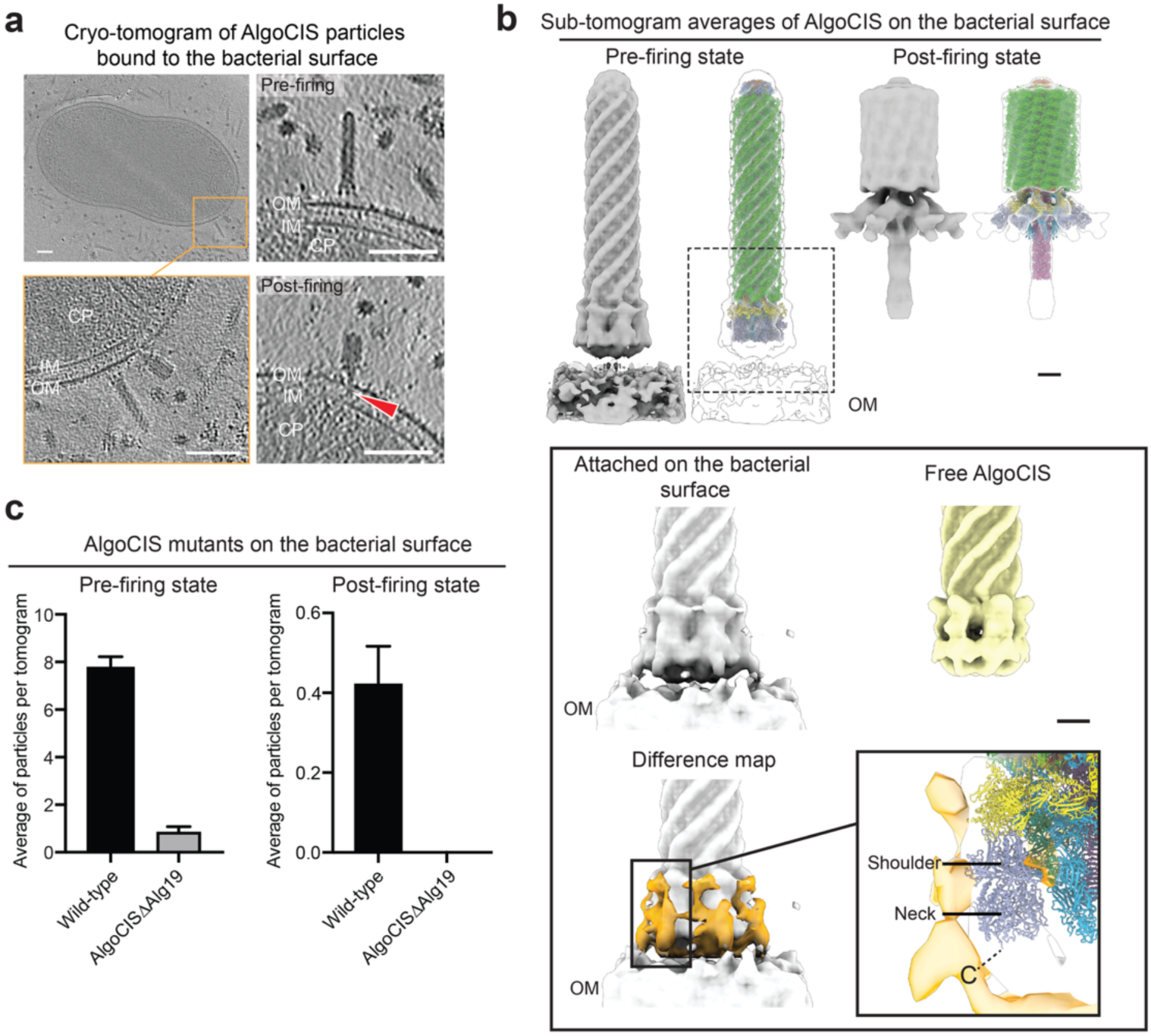
Tail-fibers mediate AlgoCIS binding to bacterial cells. **a.** Cryo-tomogram showing the pre- and post-firing AlgoCIS on the surface of *E. pacifica*. The overview is shown on the top left, the zoom-in of the boxed area (orange) is shown on the bottom left. Further zoom-ins of AlgoCIS particles on the bacterial surface are shown on the top right (pre-firing) and bottom right (post-firing). Note that the inner tube of AlgoCIS penetrates the bacterial outer membrane with the proximal end in the bacterial periplasm upon firing (red arrowhead). Shown are projections of 18 nm thick slices. OM: outer membrane; IM: inner membrane; CP: cytoplasm. Bars: 100 nm. **b.** Sub-tomogram averages of the pre- and post-firing AlgoCIS on the bacterial surface, with the structural docking of cryoEM structures. Zoom-in of the distal end of the pre-firing AlgoCIS on the bacterial surface is shown on the bottom. Note that there are additional densities (orange) around the C-terminus of the neck domain in Alg19 when compared to the free AlgoCIS. The structural components are color-coded as in Fig. 1a. The bacterial outer membrane in the post-firing AlgoCIS was masked out during data processing. Bars: 10 nm. **c.** Quantification of the pre- (left) and post-firing (right) AlgoCIS mutants on the surface of *E. pacifica* in cryo-tomograms (n = 112 in wild-type; n = 24 in AlgoCISΔAlg19). Plotted values show the mean ± SD.

We then performed sub-tomogram averaging analysis for these pre-/post-firing particles that were bound to the bacterial cell surface, showing an overall agreement with the cryoEM structures of purified AlgoCIS (Fig. 3b). For AlgoCIS in the pre-firing state; however, additional densities were detected in surface-bound complexes, when compared to the cryoEM structure of purified particles. These densities may correspond to the C-terminal part of Alg19 protein in tail-fibers, which becomes structured in bound particles but is rather flexible in free AlgoCIS (Extended Data Fig. 7c). The notion that the Alg19 tail-fiber is critical for cell surface binding was further corroborated by the fact that a tail-fiber knockout mutant (AlgoCISΔAlg19) was rarely seen bound to cells in cryo-tomograms (Fig. 3c).

Having established the binding of cells via tail-fibers, we then explored possible additional roles of Alg19 in triggering firing of AlgoCIS. First, we found that in the absence of Alg19, the mutant particles (AlgoCISΔAlg19) were more sensitive to pH-triggered firing compared to the wild-type (Extended Data Fig. 7d). This is consistent with the bridging binding profile of the tail-fibers to the baseplate wedges, suggesting that the tail-fiber may stabilize AlgoCIS in the pre-firing state. Second, we tested whether tail-fiber binding alone would be sufficient to trigger firing of AlgoCIS. We therefore purified AlgoCIS particles with a C-terminal StrepII-tag on Alg19 (AlgoCIS-Alg19^StrepII^), mixed them with a Strep-Tactin resin, and performed a pulldown assay. AlgoCIS-Alg19^StrepII^ could indeed bind to and be eluted from the resin; however, all particles remained in the pre-firing state after elution (Extended Data Fig. 7e).

Altogether, these data suggest that AlgoCIS binds to bacterial cells via tail-fibers, that Alg19 may provide a stabilizing checkpoint for firing, and that AlgoCIS may require an as yet unidentified secondary receptor to trigger firing after the binding of the tail-fibers.

### Intermediate structures reveal a stepwise firing mechanism

Having determined the start- and end-points of firing, we then asked the question whether we could generate first insight into the transitions between these states. The speed of firing, however, has so far prevented the field from resolving potential structural intermediates. Based on our structural knowledge and the genetic tractability of the system, we set out to capture different intermediate states and visualize them.

First, we generated a non-contractile sheath mutant by introducing additional residues in the N-terminus of the sheath protein (NC-AlgoCIS), similar to previously reported mutants of other CIS^23,24,38^ (Extended Data Fig. 8a). We then purified particles and performed a low-pH treatment to trigger firing. We identified a subtle intermediate (state-II) by plunge-freezing the sample and solving a cryoEM structure (Extended Data Fig. 8b-d). This state-II shows only slightly outward-tilted tail-fibers (∼5°) and the cage that was only partially open (∼10°) (Fig. 4 and Extended Data Fig. 8e-f). Excitingly, negative-stain EM analysis of the same sample discovered additional two distinct intermediate structures, which we called “state-III” (68% abundance) and “state-V” (20% abundance) (Extended Data Fig. 9a). State-III and state-V both have extended sheath and outward-tilted tail-fibers; however, they differ in the conformations of the iris (narrow *vs.* expanded) and the cage (closed *vs.* open) (Fig. 4 and Extended Data Fig. 9b). This finding suggested that the signal for triggering firing may travel through an interaction between the cage and the iris.

**Figure 4.**
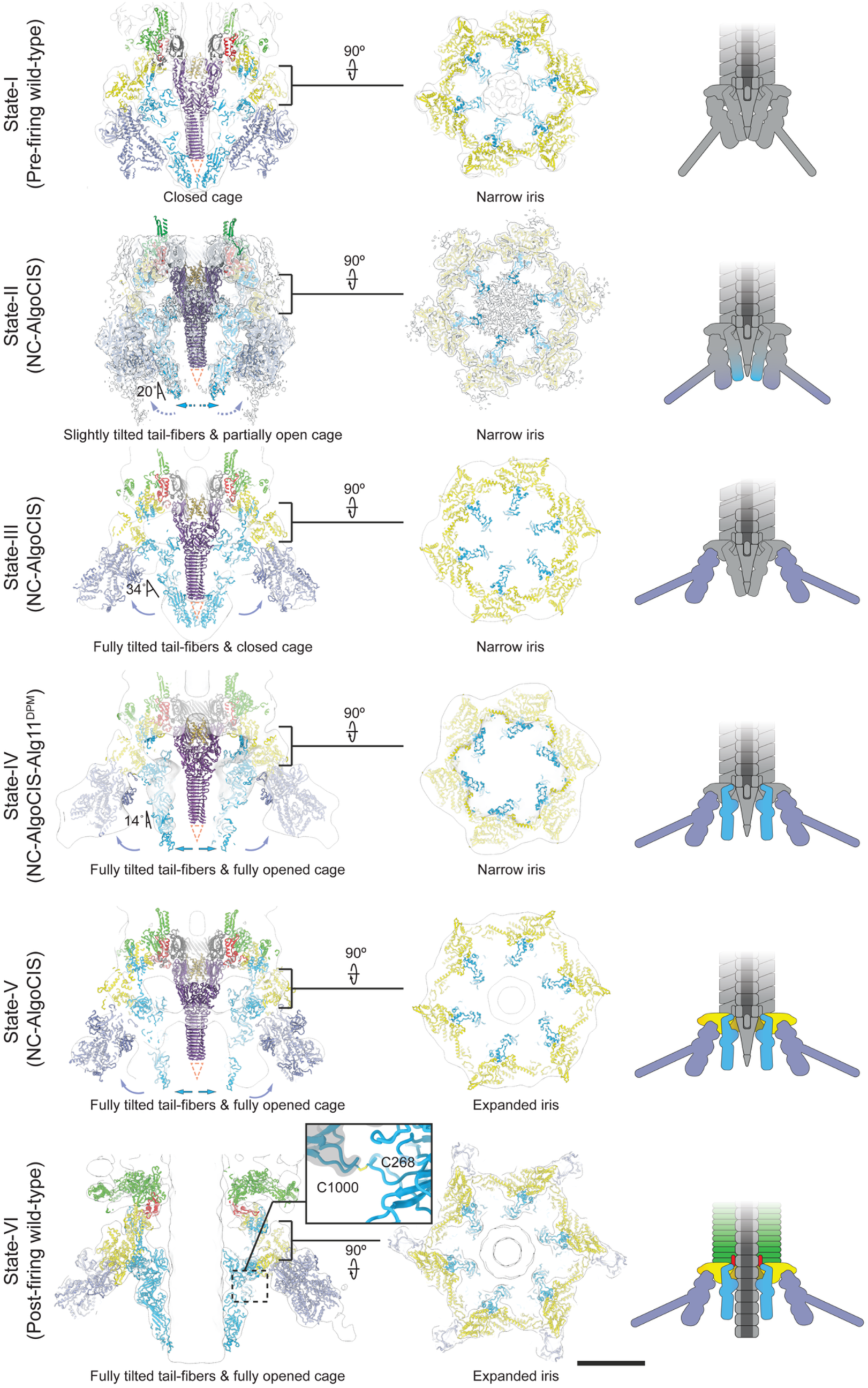
Structural snapshots of AlgoCIS intermediate states. Structures of different intermediate states (left column: central sliced view; middle column: top view of the iris; right column: schematic representation) were captured by genetic engineering (non-contractile sheath mutant, NC-AlgoCIS; disulfide bond mutant, Alg11^DPM^) and structure determination by single particle EM analysis (states-III to V: negative stain EM; state-II: cryoEM). Pre-firing (state-I) and post-firing (state-VI) structures are also shown. The position of the disulfide bond connecting the iris (shadowed by grey) and the upper part of the cage in ‘state-VI’ is boxed and zoomed in. The density maps of the pre- and post-firing structures (state-I and state-VI) were lowpass-filtered to 15 Å. Structural subunits are color-coded as in Fig. 1a. The angles between the cage and the vertical axis of the complex are indicated, structural movements are shown by arrows (slightly opened cage: dashed blue; open cage: blue; slightly tilted tail-fiber: dashed dark gray; tilted tail-fiber: dark gray). The position of the PAAR-like protein Alg10 is represented by dashed triangles. Bar: 10 nm.

To explore this hypothesis, we identified a disulfide bond (C268-C1000) in Alg11 connecting the cage to the iris (Fig. 4) and we generated a mutant (AlgoCIS-Alg11^C268A/C1000A^, hereafter referred to as AlgoCIS-Alg11^DPM^) with a defect to form this specific disulfide bond. Intriguingly, AlgoCIS-Alg11^DPM^ was not able to fire when bound to the bacterial surface (Fig. Extended Data Fig. 9c). To explore whether we could use this finding to capture a further intermediate, we then generated a double mutant, featuring non-contractile sheath and the defect to form the critical disulfide bond (NC-AlgoCIS-Alg11^DPM^). We purified particles and treated the sample with low-pH. Negative-stain EM analysis discovered yet another distinct intermediate structure (state-IV). This ‘state-IV’ shows extended sheath, a narrow iris, outward-tilted tail-fibers, and an open cage (Fig. 4 and Extended Data Fig. 9b). We hypothesized that the in total six distinct structures can be arranged in a timeline to represent a potential stepwise firing mechanism, which will be discussed below.

## Discussion

Studies of CISs *in vitro* typically reveal the systems in a pre- and/or post-firing state, without particles being seen in intermediate states. On one hand, this could be due to the fact that the native cellular context of the systems (e.g. CISs bound to the target cell) are not being investigated. On the other hand, the entire process of firing may be too fast to be captured. Here we leveraged an integrative approach that combines 1) cryoET imaging of AlgoCIS on the bacterial cell surface, 2) structure-guided engineering in order to arrest AlgoCIS in intermediate states of firing, and 3) downstream structure determination of the intermediates by single particle EM. This approach allowed us to propose a model for the propagation of the signal through the different structural modules, from the tail-fiber binding to the sheath contraction.

A model for the stepwise mechanism of AlgoCIS firing is shown in Figure 5. Cryo-tomography imaging showed that the C-terminal part of the tail-fibers is highly flexible in non-attached AlgoCIS particles (Fig. 5a). Cryo-tomograms of AlgoCIS with bacterial cells further revealed that some particles were loosely bound to the surface, likely because not all tail-fibers were yet attached to their receptors (Fig. 5b). This is consistent with previous reports showing the determination of specificity to cell surface receptors via tail-fibers for other CISs^26,28^. Like for tail-fibers in bacteriophages [e.g. long tail-fibers in T4 (ref.^39^), tail-fiber in T7 (ref.^40^)], the loose binding may help to orient the particle perpendicularly, allowing all other tail-fibers to bind to the surface, as shown in the sub-tomogram average of pre-firing AlgoCIS on the cell (Fig. 5c). These data also show that the shoulder and neck domains of the tail-fibers become less dynamic upon binding to the surface (Fig. 3b). Furthermore, upon this full attachment, we propose that the outward tilt of the tail-fibers shortens the distance between the cage and the cell surface, and it also exposes a larger fraction of the cage outer surface. Interestingly, the cage comprises two potential carbohydrate binding modules^19^, which may then be accessible for a potential as yet uncharacterized secondary receptor (Fig. 5d). Subsequently, we hypothesize that the signal from the opening of the cage is then transmitted to the iris via a disulfide bond that we identified being critical for firing (Fig. 4). As a result, the iris expands, triggers a conformational change in the sheath initiator, followed by sheath contraction, which is consistent with the mechanism of other CISs and phage T4^13,15,25,31–33^ (Fig. 5e). The contraction proceeds from the proximal to the distal end of sheath-tube module. Finally, sheath contraction propels the inner tube forward, and it releases the cap adaptor at the distal end of the complex (Fig. 5f).

**Figure 5.**
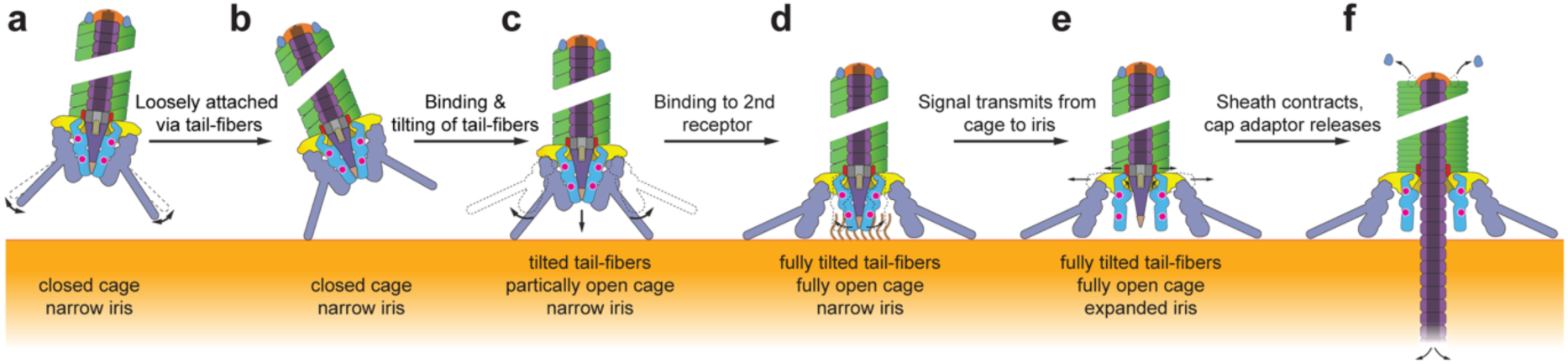
Putative stepwise AlgoCIS firing mechanism. Schemes showing a stepwise firing mechanism in AlgoCIS. The C-terminal part of the tail-fibers is flexible in non-attached AlgoCIS particles (**a**). The flexible C-terminal part of the tai-fibers can mediate the initial binding of AlgoCIS to the target surface via loose attachment (**b**). The loose binding further helps to orient the particle perpendicularly for the full attachment. The tail-fibers tilt outwards, shortening the distance between the cage and the cell surface (**c**). The full tilting of tail-fibers exposes a larger fraction of the cage outer surface. The two potential carbohydrate binding modules (magenta hexagons) then are accessible to bind an as yet uncharacterized secondary receptor (brown), resulting in a fully opened cage (**d**). The signal propagates to the iris and triggers its expansion, inducing a conformational change in the sheath initiator, followed by sheath contraction (**e**). The sheath contraction proceeds from the proximal to the distal end of sheath-tube module. Finally, sheath contraction propels the inner tube forward and releases the cap adaptor at the distal end of the complex (**f**).

In summary, our study indicates a stepwise firing mechanism of AlgoCIS that may require two sequential binding events to the cell surface via the tail-fibers and the cage, respectively. This is consistent with our finding that the binding of AlgoCIS tail-fibers to coated beads was not sufficient to trigger firing (Extended Data Fig. 7e). This aspect is distinct from the ‘checksum’ mechanism that was proposed for pyocin R2 (ref.^15^). In these cage-less CISs and also potentially in the cage-less PVCs, tail-fiber binding alone may be sufficient to trigger the firing. Interestingly, the stepwise firing mechanism of AlgoCIS proposed here shows remarkable similarities to the bacterial toxin complex (Tc) and the T4 phage, both of which require the sequential interactions with specific receptors to result in a tight binding to the target cell surface before fully functioning^11,39,41,42^. These potential checkpoints may function together to avoid non-productive firing events.

Our study will serve as a framework for understanding the firing mechanism of other CISs that have conserved structural modules and even function by a different more of action, such as representatives of the T6SS*^iv^*, tCIS, and cytoplasmic CIS. Furthermore, our innovative approach can be applied not only to these systems, but also to more distantly related CISs including the T4 phage or pyocins. Finally, the structural and mechanistic insights here will facilitate the re-engineering of AlgoCIS and related CISs as a biomedical tool for the delivery of cargo proteins of choice to specific target cell lines.

---

**Extended Data Figure 1.**
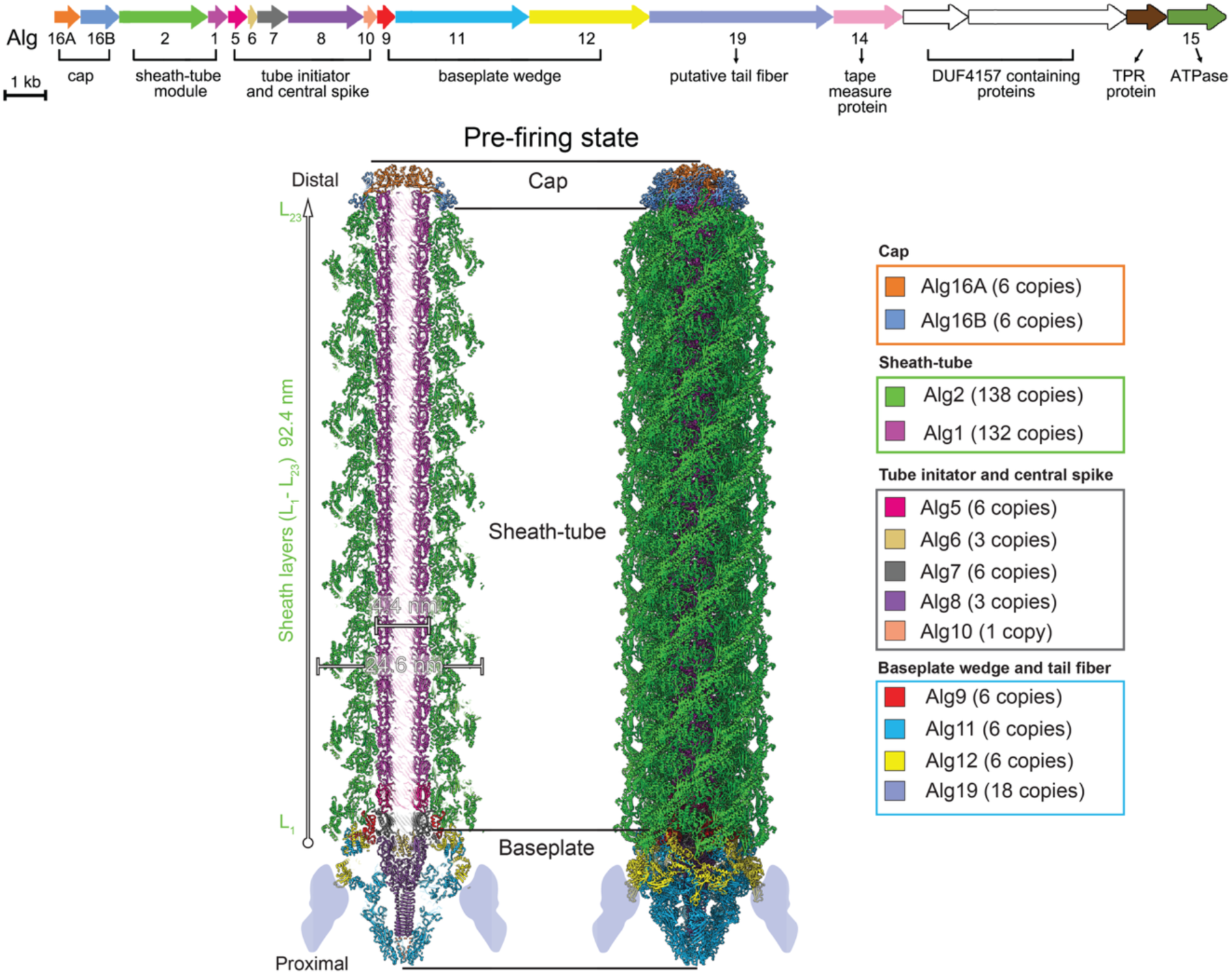
Structure of the AlgoCIS in the pre-firing state. Atomic model of AlgoCIS in the pre-firing state (left: central sliced view; right: side view), showing that AlgoCIS comprises three modules (cap, sheath-tube, and baseplate). The gene cluster of the contractile injection system in *A. machipongonensis* (AlgoCIS) is shown on the top, the genes encoding cargo proteins are colored white. Structural subunits are color-coded, while the positions of the tail-fibers (Alg19) are shadowed in dark gray. The gene cluster and the model are derived from a previous study^19^.

**Extended Data Figure 2.**
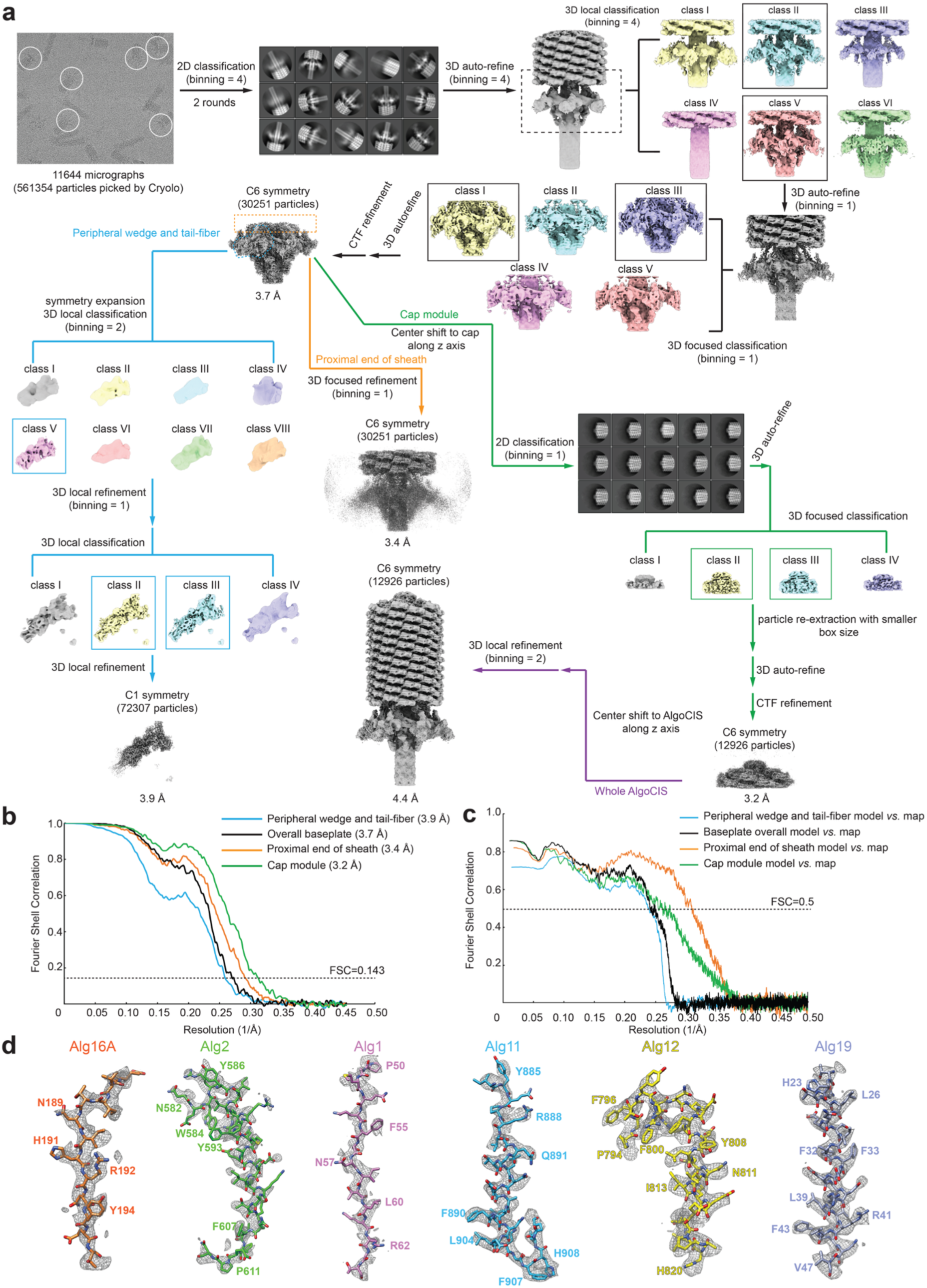
CryoEM data processing workflow of the post-firing AlgoCIS treated with guanidine-HCl. **a.** Flowcharts for the cryoEM reconstruction of different parts of the post-firing AlgoCIS treated with guanidine-HCl. See methods for details. **b-d.** Gold-standard Fourier shell correlation (FSC) (**b**) and model *vs.* map FSC (**c**) curves of the cryoEM reconstruction of different parts of the post-firing AlgoCIS treated with guanidine-HCl. The representative densities on different proteins are shown in (**d**).

**Extended Data Figure 3.**
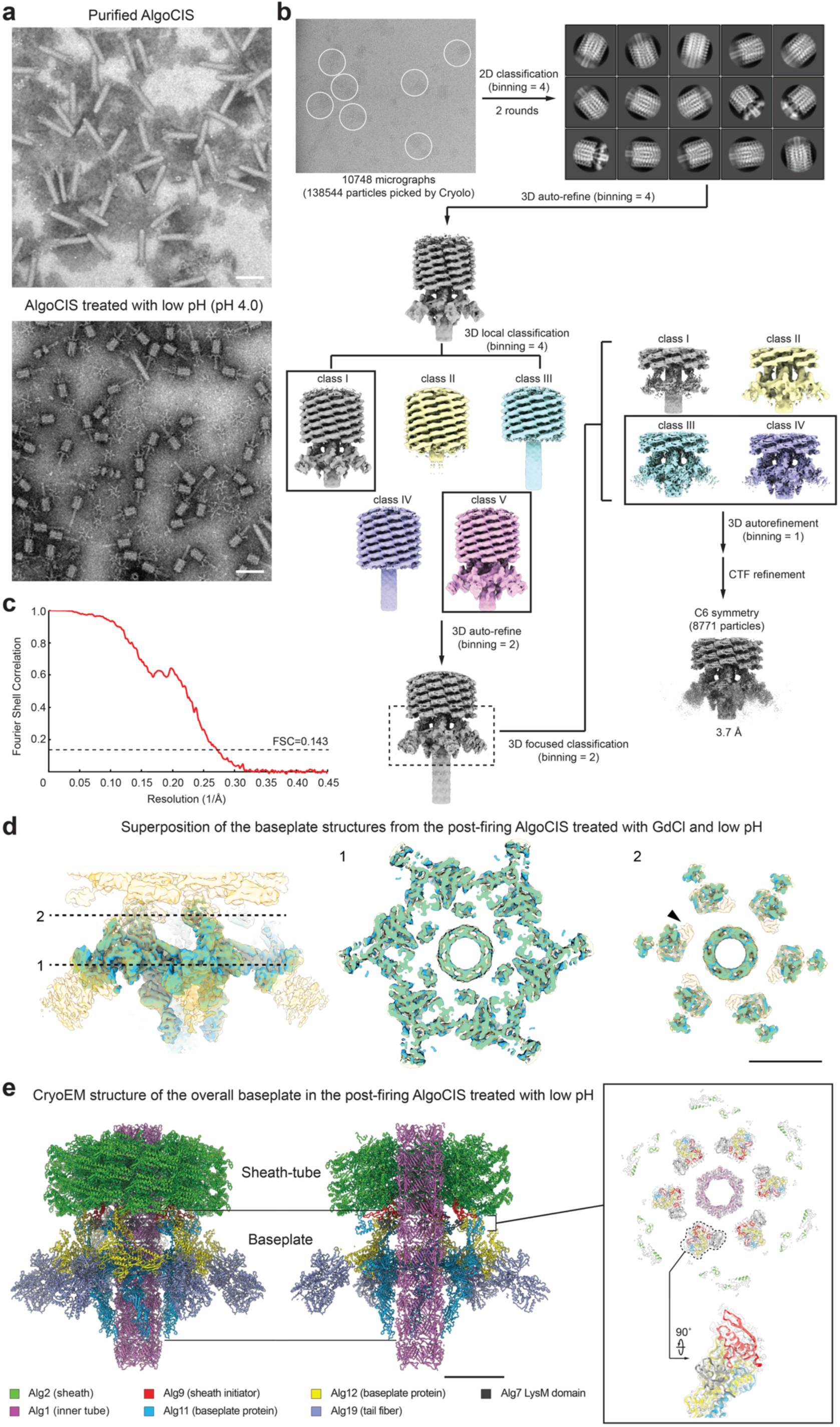
CryoEM data processing workflow of the post-firing AlgoCIS treated with low pH. **a.** Negative-stain EM images of purified AlgoCIS with/without low pH treatment, showing that low pH (the value of pH is 4.0) can trigger AlgoCIS firing. Bars: 100 nm. **b.** Flowcharts for the cryoEM reconstruction of the baseplate in the post-firing AlgoCIS treated with low pH. See methods for details. **c.** Gold-standard FSC curve of the cryoEM reconstruction of the baseplate in the post-firing AlgoCIS treated with low pH. **d.** Superposition of the baseplate structures from the post-firing AlgoCIS treated with guanidine-HCl (blue, presented at ±7.6σ) and low pH (orange, shown in transparent, presented at ±3.8σ), showing the overall structural agreement (correlation: 0.98) of the baseplate. The cross-section views at different positions are shown on the right. Both maps were lowpass-filtered to 6 Å. Note that there is one additional density (black arrowhead) observed in the structure from the sample that was treated with low pH, which is formed by LysM domain in Alg7. Bar: 10 nm. **e.** Atomic model of the baseplate in the post-firing AlgoCIS treated with low pH (left: side view; middle: central sliced view), showing the almost same structure of the baseplate as the one treated with guanidine-HCl. Cross-section view of the proximal end of the sheath is shown on the top right, where one baseplate wedge is outlined and the side view is shown on the bottom right. Note that LysM domain of Alg7 is found in the post-firing AlgoCIS treated with low pH, but is absent in the model treated with guanidine-HCl. Structural subunits are color-coded as in Fig. 1a. Bar: 10 nm.

**Extended Data Figure 4.**
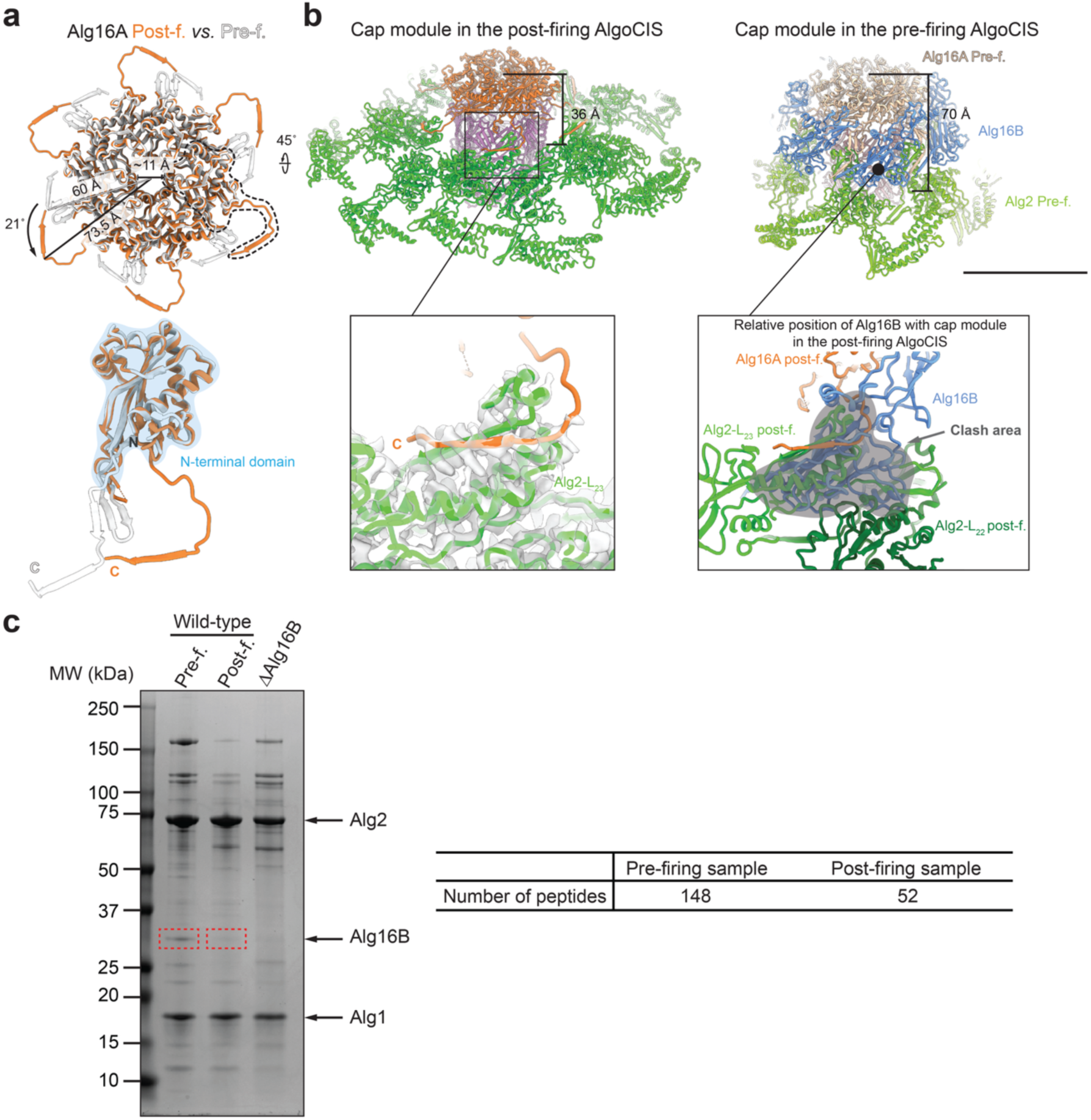
Structural re-arrangement of the cap module in AlgoCIS upon firing. **a.** Structural comparison of the cap protein Alg16A in the pre- (white) and post-firing (orange) states, showing that the C-terminus of Alg16A rotates 21° upon firing. The inner and outer diameters of six Alg16A proteins were measured. One Alg16A is outlined, and the side view is shown on the bottom, with N-terminal domain shadowed in cyan and the N- and C-termini labeled. **b.** Structural comparison of the cap module of in the post- (left) and the pre-firing (right) AlgoCIS, showing that the cap adaptor (Alg16B: blue) is absent in the post-firing AlgoCIS. The distance between the cap and the proximal sheath layer (Alg2-L_23_) decreases upon firing (contracted: 36 Å *vs.* extended: 70 Å). The contact sites between the cap and the distal sheath layer are zoomed-in and shown on the bottom. Note that severe clashes (shadowed in grey) are observed between the cap adaptor (in the pre-firing state) and the distal sheath layer (in the post-firing state) when superpositioned based on the N-terminal domain of the cap, suggesting that the cap adaptor is extruded upon firing. Structural subunits are color-coded as Fig. 1a, the cap protein (Alg16A) and the distal sheath layer in the pre-firing state is colored brown and light green. The second distal sheath layer (Alg2-L_22_) in the post-firing state is colored dark green. The EM map is shown transparent. Bar: 10 nm. **c.** SDS-PAGE and mass spectrum analysis detected a lower amount of the cap adaptor protein (Alg16B) in the purified post-firing sample. Left: SDS-PAGE gel of purified AlgoCIS in the pre- and post-firing states (lane 1-2), AlgoCISΔAlg16B (lane 3) was a control. The bands of the sheath (Alg1), inner tube (Alg2), and the cap adaptor protein (Alg16B) are indicated, where Alg1 and Alg2 are regarded as loading controls. The mass spectrum analysis on the Alg16B (dashed red boxes) is shown on the right. The number of the detected Alg16B peptide in both samples are indicated.

**Extended Data Figure 5.**
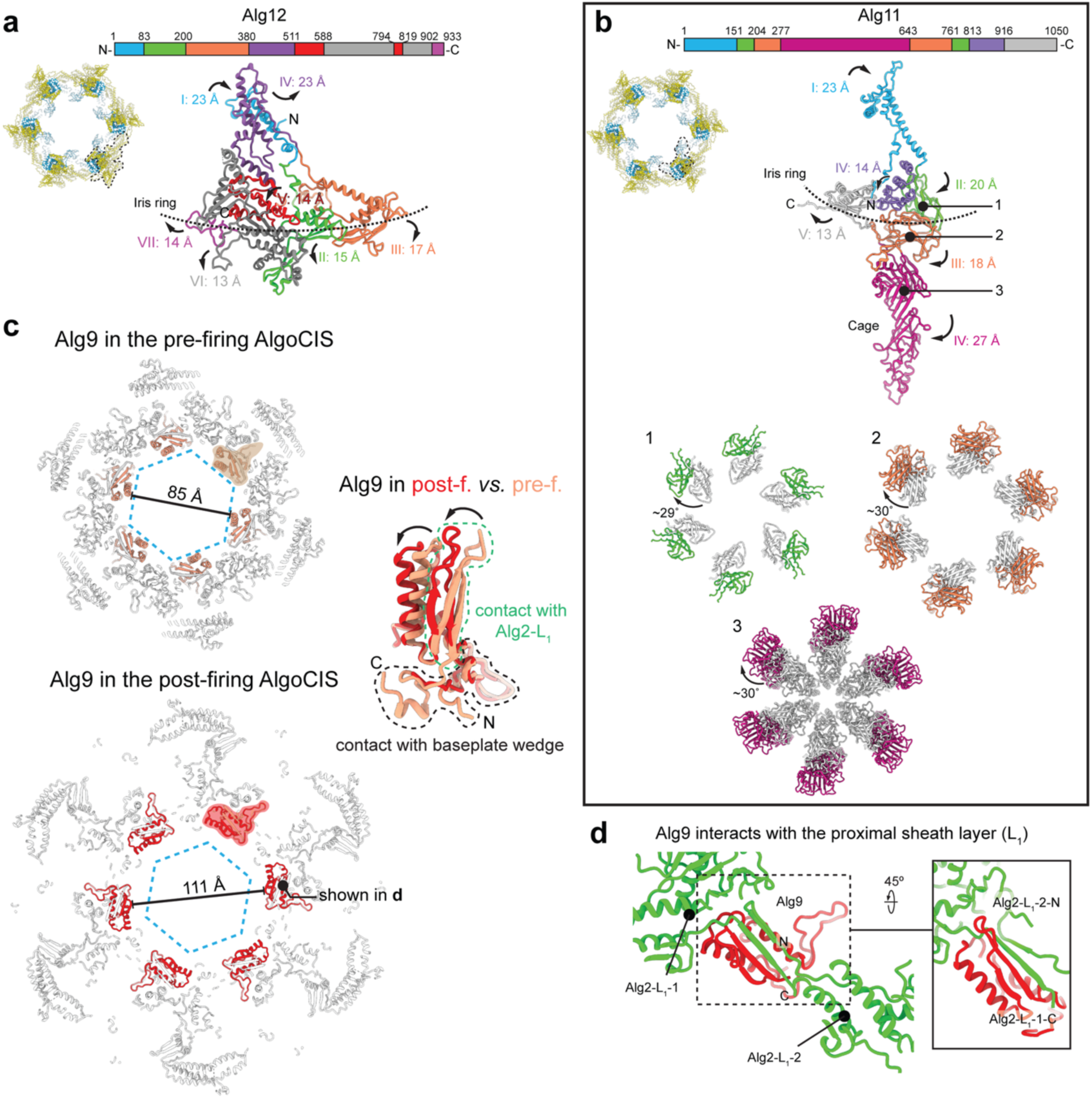
Structural re-arrangement of the baseplate in AlgoCIS upon firing. **a.** Individual domains of Alg12 have different degrees of outward movement upon firing. Left: Cross-section view of the iris (color-coded as in Fig. 1a), one Alg12 protein is outlined. Right: Domain organization of Alg12 (top) and the structure of the Alg12 protein in the post-firing state (bottom). Domains are color-coded, the position of the iris is represented by dashed line. Movements of individual domains are indicated by arrows, where the values were measured based on the shifts of the centers of each domain in the pre- and post-firing states. **b.** Individual domains of Alg11 have different degrees of outward movement upon firing. Top: Cross-section view of the iris (shown as panel **a**), where one Alg11 protein is outlined. Middle: Domain organization of Alg11 (top) and the structure of the Alg11 protein in the post-firing state (bottom). Style is shown same as in (**a**). The movements of individual domains are indicated by arrows and are measured. Bottom: Cross-section views on different domains of Alg11 showing that the domain II (1), domain III (2), and domain IV (3, cage) have both rotation (∼30°, indicated by arrows) and outward expansion upon firing. The Alg11 proteins in the pre-firing state are colored white. **c.** Structural comparison of the sheath initiator protein (Alg9) in the pre- (top-left, coral) and post-firing (bottom-left, red) states, showing that Alg9 has outward movement upon firing. The diameters of Alg9 rings are labeled, the positions of the inner tube are presented by the dashed hexagon (blue). One Alg9 protein in the pre- and post-firing states are shadowed and the structural comparison is shown on the right. The structural re-arrangements are indicated by arrows, the regions that interact with the proximal sheath layer or the baseplate Alg11/12 heterodimer are outlined by green or black dashed line, respectively. **d.** Alg9 interacts with two adjacent sheath proteins via hand-shake interactions in the post-firing state. Structural subunits are color-coded as in Fig. 1a.

**Extended Data Figure 6.**
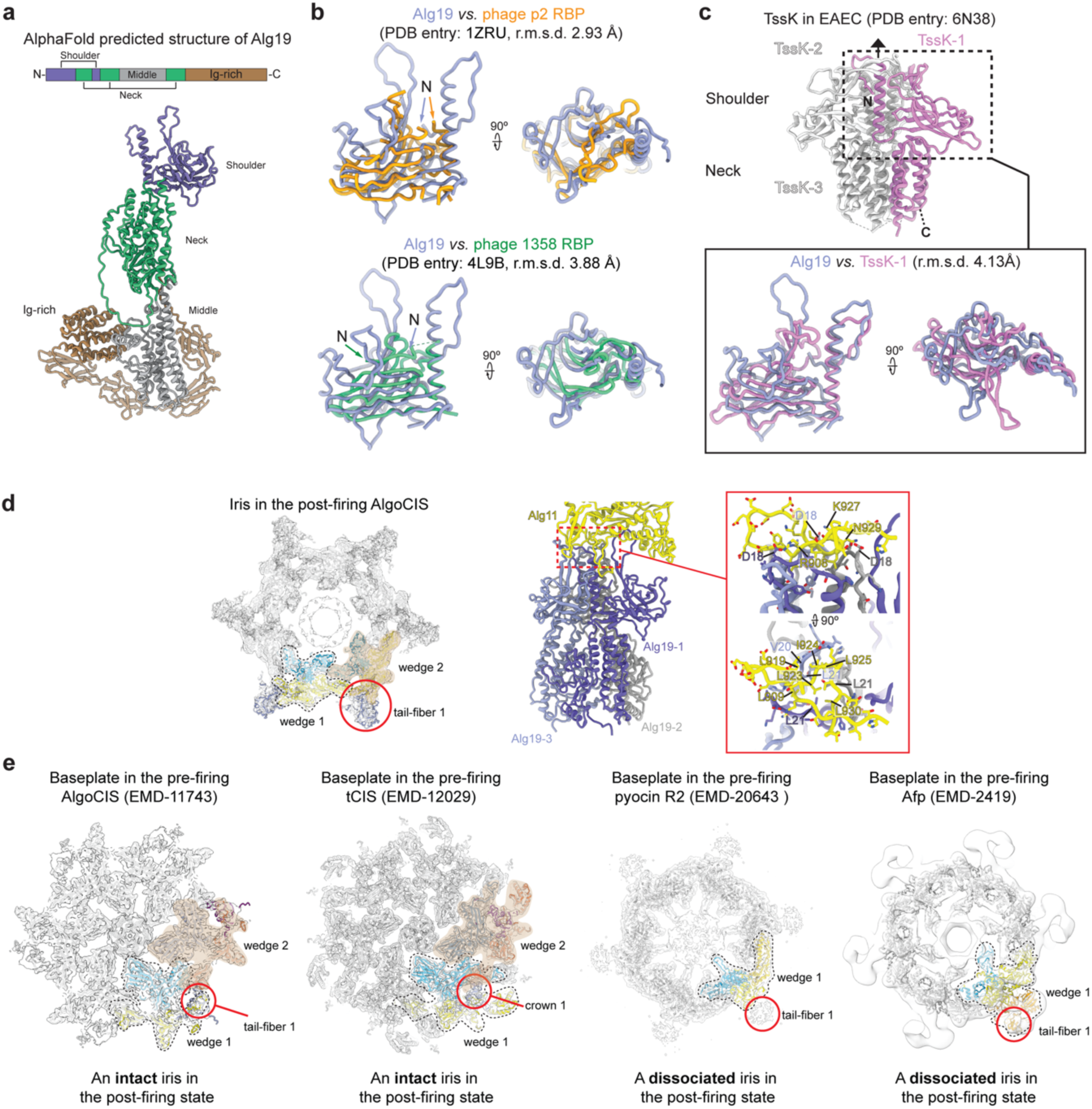
Tail-fibers bind to the baseplate. **a.** Alg19 is predicted to have four domains. Domain organization of Alg19 is shown on the top, and the Alphafold predicted structure is shown on the bottom and color-coded. **b.** The shoulder domain of Alg19 has a similar structure as some phage receptor binding proteins (RBPs) (top: lactococcal phage p2 RBP, PDB-1ZRU, orange; bottom: lactococcal phage 1358 RBP, PDB-4L9B, green). The overall RMSD values are indicated. **c.** The shoulder domain of Alg19 has a similar structure as the shoulder part of the T6SS TssK in enteroaggregative *Escherichia coli* (EAEC). Structure of trimeric TssK (PDB-6N38) is shown on the top, one protomer is colored pink. The shoulder domain of TssK is highlighted by dashed box, while the structural superposition with the shoulder domain of Alg19 is shown on the bottom. The overall RMSD value is indicated. **d.** One tail-fiber connects two adjacent baseplate wedges in AlgoCIS. Left: cross-section view of the baseplate in the post-firing AlgoCIS. One baseplate wedge is outlined and its structural subunits (wedge 1) are color-coded as in Fig. 1a. The adjacent baseplate wedge (wedge 2) is shadowed in brown and structural subunits are colored (Alg11: coral; Alg12: dark blue; tail-fiber: magenta). The tail-fiber binding site is highlighted by red circle. EM maps are shown in transparent. Right: side view of one tail-fiber binding to the baseplate. Each protomer of tail-fibers and Alg11 are colored. The binding sites are boxed and the zoom-in is shown on the right. The residues participating in contacts are presented with side chain and are labeled. Note that both hydrophobic interaction and salt bridges mediate the interactions. **e.** The tail-fibers have different binding profiles to the baseplate among CISs. Shown are cross-section views of the baseplate structure from different CISs in the pre-firing state (left to right: AlgoCIS, tCIS, pyocin R2, and Afp). The structures are shown in the same style as panel (**d**). Note that one tail-fiber binds to an individual baseplate wedge in pyocin R2 and Afp, which has a dissociated iris upon firing, while the tail-fiber/crown connects two adjacent baseplate wedges in AlgoCIS/tCIS that keeps an intact iris upon firing.

**Extended Data Figure 7.**
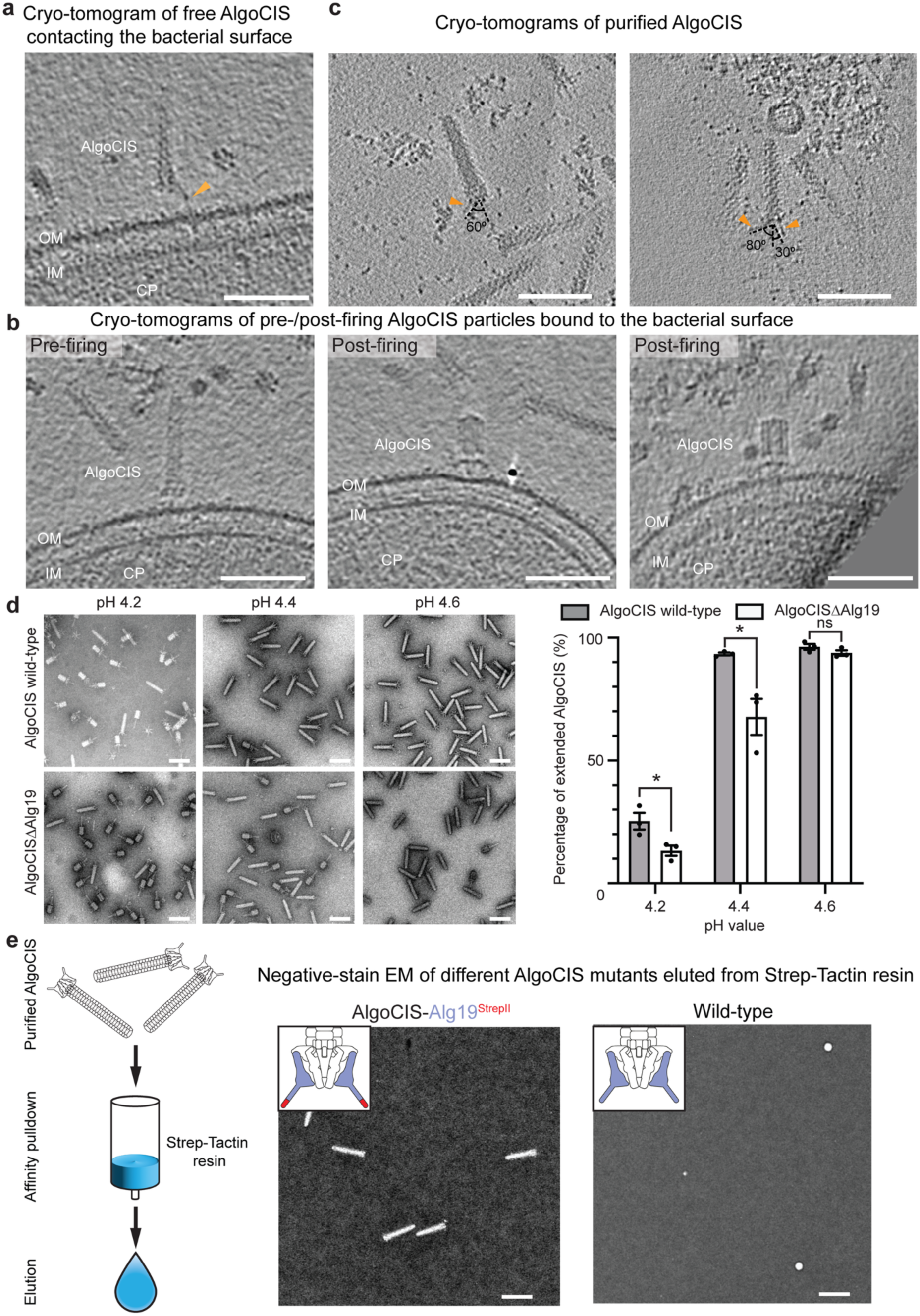
Tail-fibers mediate the AlgoCIS attachment to bacterial surface a-b. Additional examples in cryo-tomograms of co-incubation of AlgoCIS wild-type with *E. pacifica*. A pre-firing AlgoCIS particle contacts the bacterial surface via its flexible tail-fiber (**a**, orange arrowhead). A pre-firing AlgoCIS perpendicularly attaches to the bacterial surface (**b**, left panel). AlgoCIS fired on the bacterial surface (**b**, middle and right panels). Shown are projections of 18 nm thick slices. Bars: 100 nm. OM: outer membrane; IM: inner membrane; CP: cytoplasm. **c.** Cryo-tomograms showing that the C-terminal part of the tail-fiber is highly flexible in non-attached AlgoCIS. Shown are projections of 10.8 nm. The tail-fibers are indicated with orange arrowheads, and the angles are measured. Bars: 100 nm. **d.** AlgoCIS is more sensitive to low pH treatment when knocking out the tail-fibers. Shown are representative negative-stain EM images of wild-type and the tail-fiber knockout mutant (AlgoCISΔAlg19) upon treatments with different pH values. Quantification is shown on the right. Note that some AlgoCISΔAlg19 contract in the buffer with pH value of 4.4, but almost all particles of wild-type remain in the pre-firing state. Plotted values show the mean ± SD from three independent experiments. Bars: 100 nm. **e.** Schematic (left) and negative-stain EM images (right) showing that the purified AlgoCIS fused with C-terminal StrepII-tag (AlgoCIS-Alg19^StrepII^) remains in the pre-firing state after StrepII-tag pulldown, indicating that the binding of the tail-fiber to a receptor is insufficient to trigger the firing. The wild-type is regarded as negative control. Bars: 100 nm.

**Extended Data Figure 8.**
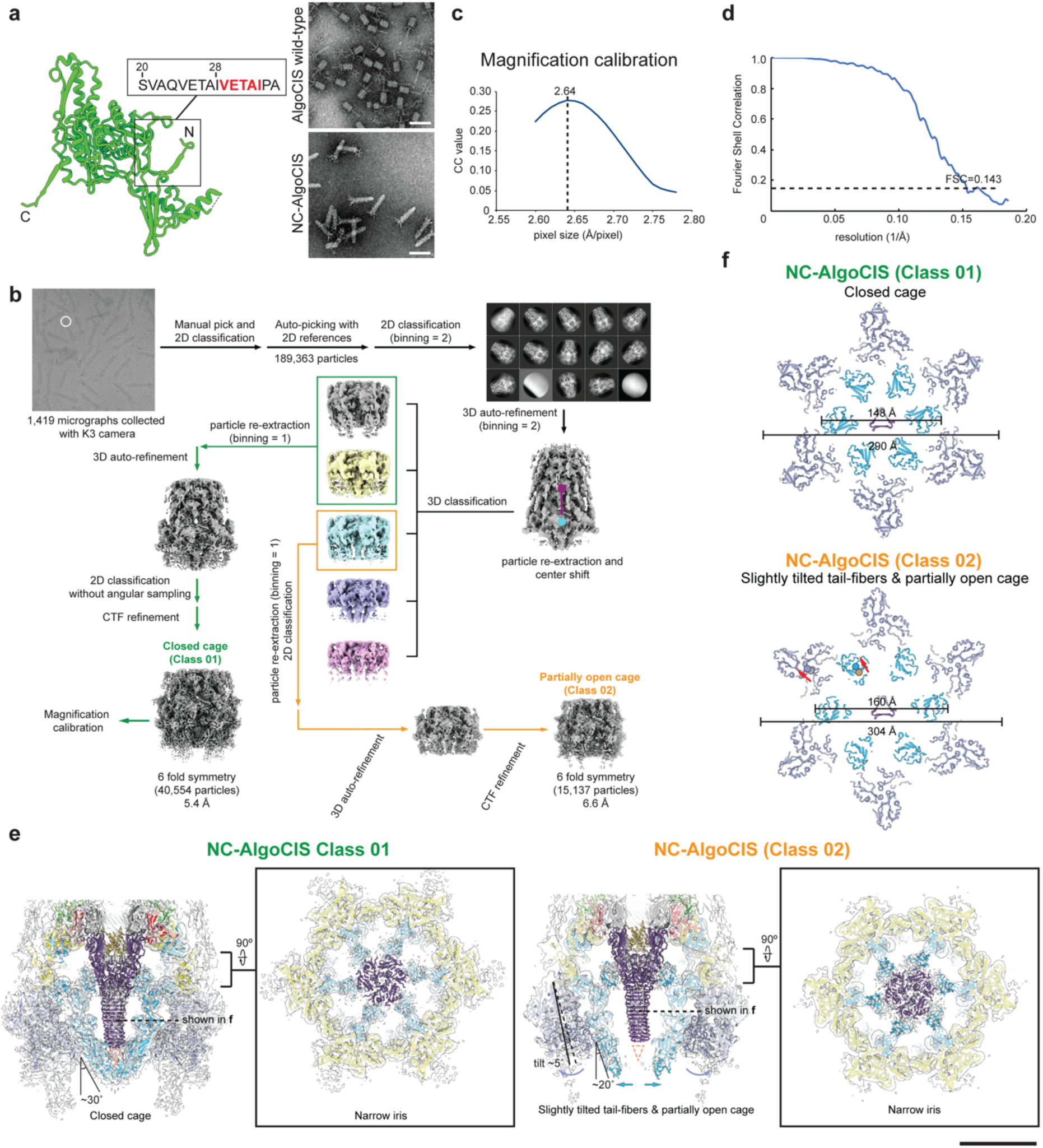
CryoEM analyses of an AlgoCIS intermediate state. **a.** Re-engineering of the non-contractile AlgoCIS (NC-AlgoCIS) mutant by introducing five additional amino acid residues (red) into the N-terminal part of sheath protein Alg2. Representative negative-stain EM images are shown on the right. Note that NC-AlgoCIS cannot contract upon low pH (pH value of 4.0) treatment, while all wild-type particles fire in the same condition. Bars: 100 nm. **b.** Flowcharts for the cryoEM reconstruction of the non-contractile AlgoCIS mutant (NC-AlgoCIS) treated with low pH. See methods for details. Note that two classes of NC-AlgoCIS were determined from the dataset: one is in the pre-firing state and has a closed cage (Class 01); the other is in the pre-firing state but has slightly tilted tail-fibers and a partially opened cage (Class 02). **c.** Plot of magnification calibration on the dataset of NC-AlgoCIS treated with low pH. See methods for details. **d.** Gold-standard FSC curve of the cryoEM reconstruction of the baseplate with slightly tilted tail-fibers and a partially opened cage in NC-AlgoCIS (Class 02) treated with low pH. **e-f.** NC-AlgoCIS with slightly tilted tail-fibers has a partially opened cage upon low pH treatment. Shown are the two classes of NC-AlgoCIS treated with low pH. The central sliced views of the overall baseplate structure (left) and the cross-section view of the iris (right) are shown in panel **e**. Structural re-arrangements on the cage and tail-fibers are indicated by arrows (blue: cage; dark gray: tail-fibers). The angles between the cage and the perpendicular axis were measured. The outward tilted angle is labeled, where the axis of the tail-fibers with/without outward tilting is presented with a solid or dashed line, respectively. The cross-section view on the tail-fibers is shown in panel **f**. The outer-diameters of the cage and the tail-fibers were measured. The shifts of the centers of the cage (Class 01: orange; Class 02: blue) and the tail-fibers (Class 01: brown; Class 02: dark grey) are indicated by red arrows. Bar: 10 nm.

**Extended Data Figure 9.**
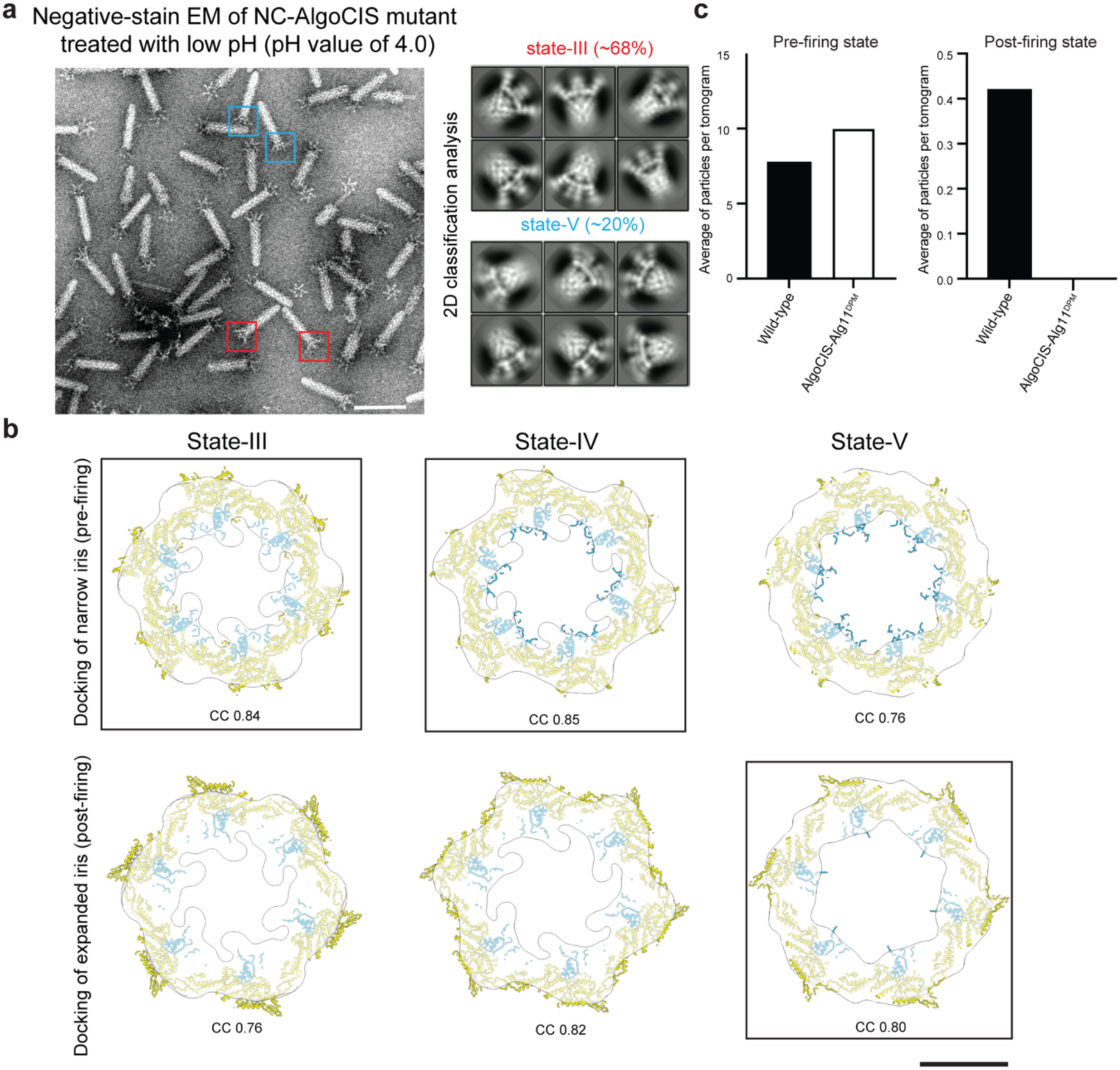
Different intermediates are observed in NC-AlgoCIS treated with low pH. **a.** Two different intermediate structures were observed when NC-AlgoCIS was treated with low pH and imaged by negative-stain EM. Left: one representative negative-stain EM image of NC-AlgoCIS upon low pH treatment, where two intermediates are highlighted by colored boxes (state-III: red; state-V: blue). Right: 2D classes of two intermediates (state-III: top; state-V: bottom), the percentages of each state are indicated. Bar: 100 nm. **b.** Structural fittings of the pre-(top)/post-firing (bottom) iris into reconstructed maps of intermediate structures, showing that the iris of state-III and state-IV are in the pre-firing state, and the iris of state-V is in the post-firing state. Correlation coefficient (CC) values are stated. Iris structures with best CC value are boxed. See methods for details. Bar: 10 nm. **c.** Quantification of the pre- (left)/post-firing (right) AlgoCIS mutants on the surface of *E. pacifica* in cryo-tomograms (n = 112 in wild-type; n = 24 in AlgoCIS-Alg11^DPM^). Note that no firing events are observed in *E. pacifica* co-incubated with AlgoCIS-Alg11^DPM^. The dataset of wild-type is the same as shown in Fig. 3c.

## Methods

### Growth and handling of Bacterial strains

*Algoriphagus machipongonesis* PR1 and *Echinicola pacifica* bacteria were grown in fresh marine broth (MB) media (Condalab) and grown at 30 °C and 200 r.p.m. for 2 days unless otherwise stated.

### Generation of mutant strains

All bacterial strains, plasmids, and primer sequences used are listed in Extended Data Table 2. All mutant strains were created according to previously published protocols^19^. Briefly, the bacteria were mated with *E. coli* SM10 (λpir) that harbored the pCHIP3 plasmid containing the mutation of interest. Plasmid integration was selected with agar plates containing 50 µg/ml Erythromycin, followed by a counter-selection on MB plates containing 10 mM 4-chloro-DL-phenylalanie. Correct mutants were confirmed via PCR and sanger sequencing.

### AlgoCIS purifications

Different AlgoCIS mutants were purified following the previously report^19^. Briefly, the bacterial culture of *A. machipongonesis* mutants was grown in 1 L of MB media for 48 hrs at 30 °C and 200 r.p.m.. The bacteria were harvested via centrifugation at 9000 g for 20 min at 4 °C. The pellet was then resuspended in 20 ml of residual media and the lysis reagents [the final concentration of 1% (v/v) Triton-X100, 0.5 × CellLytic B (Sigma-Aldrich), 200 μg/ml lysozyme, 50 μg/ml DNaseI] were added. The bacteria were lysed by incubation at 37 °C for 30 min and the cell debris was removed via centrifugation at 21,000 g for 20 min. The supernatant was then subjected to ultra-centrifugation at 150,000 g and 4 °C for 1 hour, where samples were split and laid onto a 1 ml sucrose cushion [20 mM Tris pH 8.0, 150 mM NaCl, 50 mM EDTA, 1% Triton-X100, 50% (w/v) sucrose] for each centrifuge tube. After spinning, the cushion was then taken along with ∼1 ml the overlying liquid and was centrifuged at 21,000 g for 15 min to remove any residual cellular debris. The supernatant was subjected to a second round of ultra-centrifugation without a sucrose cushion. The resulting pellets were then soaked in TS-buffer (20 mM Tris pH 7.5 and 150 mM NaCl) overnight at 4 °C, then were resuspended. Crude samples were further purified through a 10-50% (w/v) sucrose gradient at 100,000 g and 4 °C for 1 hour using SW55 Ti rotor. The fractions containing AlgoCIS particles were pooled, diluted with TS-buffer, and passed through 0.1 µm-pore filter twice. The sample was further concentrated through a third round of ultra-centrifugation (150,000 g and 4 °C for 1 hour) and the pellets were resuspended in TS-buffer. The purified particles were stored at 4 °C until used.

To prepare the contracted AlgoCIS, the purified particles were treated either with 2 M guanidine-HCl (GdCl) at room temperature for 30 min or low pH buffer (0.1 M sodium citrate buffer with a pH value of 4.0-4.8) at room temperature for 2 hrs. For the samples treated with low pH, the pH was adjusted back after incubation using 1 M Tris (pH 8.0). For the GdCl-treated samples, the residual GdCl was removed through dialysis against TS-buffer using Slide-A-Lyzer MINI dialysis devices (Thermo Fisher).

### Pulldown assays of AlgoCIS

The purified AlgoCIS wild-type or AlgoCIS-Alg19^StrepII^ was applied to StrepII-tag affinity purification (Strep-Tactin resins, IBA) and the particles were eluted after several washing steps using TS-buffer containing 5 mM desthiobiotin (Sigma). The eluted samples were stained using 1% phosphotungstic acid (PTA, pH 7.0) and imaged on a Morgani TEM (Thermo Fisher).

### Mass spectrometry analysis

To analyze the protein level of the cap adaptor protein in the extended and contracted AlgoCIS, the contracted AlgoCIS samples were prepared as above and then applied to another round of ultra-centrifuge to pellet down the contracted particles. The protein samples (the extended and contracted AlgoCIS, and the purified AlgoCISΔAlg16B) were used for SDS-PAGE analysis. The bands of Alg16B in the extended and contracted AlgoCIS were cut and sent to the Functional Genomics Center Zürich (FGCZ), which performed mass spectrometry and the data analysis.

The gel bands were cut in small pieces and washed with 100 mM NH_4_HCO_3_/50% acetonitrile and 100% acetonitrile, respectively. The samples were then performed protein digestion using trypsin (5 ng/µl in 10 mM Tris pH 8.2 and 2 mM CaCl_2_). The peptides from the supernatants were extracted (0.1% PFA, 50% acetonitrile), dried, and then dissolved in 3% aqueous acetonitrile with 0.1% formic acid. The samples were then transferred to the autosampler vials for liquid chromatography-mass spectrometry analysis (LC-MS/MS). 10 µl of each sample were injected on a M-class UPLC coupled to a Q-Exactive mass spectrometer (Thermo Fisher). The acquired MS data were imported into PEAKS Studio (Bioinformatic Solutions) and the data were searched against the *A. machipongonensis* database. The results were visualized by Scaffold software.

### CryoEM sample preparation

All cryoEM and cryoET samples were prepared using a Vitrobot Mark IV (Thermo Fisher). For the contracted AlgoCIS treated with GdCl or low pH, 4 µl of sample was applied onto Quantifoil copper grids (R2/1, 300 mesh) coated with 1 nm thickness of continuous carbon layer. The samples were plunge-frozen into liquid ethane/propane [37% (v/v) ethane]^43^ after blotted by filter papers. The cryoEM sample of NC-AlgoCIS treated with low pH was prepared similarly as above, where an extra 150 mM NaCl was added into the samples before plunge-freezing.

### Co-incubation of *E. pacifica* with AlgoCIS

For the co-incubation of *E. pacifica* with different AlgoCIS mutants, the bacterial culture of *E. pacifica* was growth in marine broth at 30 °C and 200 r.p.m. for 2 days. The bacteria were then harvested and washed with AKCGM3/AKSWC media [1.5% AKCGM3 (artificial seawater with cereal grass) and 1.5% AKSWC (artificial seawater complete)]. The bacterial suspension was adjusted to OD_600_ value of 3.0 with AKCGM3/AKSWC media, and was then mixed with the purified AlgoCIS mutant (wild-type, AlgoCISΔAlg19, AlgoCIS-Alg11^DPM^) at the ratio of 5:1. The mixture was incubated at 30 °C for 1 hour. The residual free AlgoCIS particles were removed by centrifuge at 5000 g for 5 min and the pellets were resuspended by AKCGM3/AKSWC media to bacterial OD_600_ value of 2.0.

To vitrify the co-incubation of *E. pacifica* with different AlgoCIS mutants, the samples were mixed with 10 nm Protein A-coated colloidal gold particles at a ratio of 1:5. 4 µl of sample was applied onto EM grids (R2/2, 200 mesh, Quantifoil), blotted from the backside (using a Teflon sheet on one side), and then plunged into liquid ethane/propane.

### pH stability analysis of AlgoCIS mutants

The stability of AlgoCIS mutants against different pH buffers were estimated based on the ratio of the particles in the extended and contracted state. Briefly, the purified AlgoCIS mutants (wild-type and AlgoCISΔAlg19) were treated with low pH buffer (0.1 M sodium citrate buffer with a pH value of 4.0-4.8) at room temperature for 2 hrs. The pH of sample was then adjusted back after incubation using 1 M Tris (pH 8.0). The samples were then stained by 1% phosphotungstic acid (PTA, pH 7.0) and imaged on a Morgani TEM.

To analyze the ratio of the particles in the extended and contracted states, AlgoCIS in both states were picked automatically from negative-stain EM dataset via *cryolo*^44^. The picked particles were then manually checked and used for the quantification analysis. The plots were prepared using Prism v10.

### CryoET data collection and cryo-tomogram reconstruction

All cryoET dataset were collected on a Titan Krios transmission electron microscope (Thermo Fisher) operating at 300 kV and equipped with BioContinuum imaging filter and K3 direct electron detector (Gatan). Tilt series were collected using SerialEM^45^.

For the co-incubation of *E. pacifica* with different AlgoCIS mutants, tilt series were collected in a dose-symmetric scheme with an angular range from +60° to -60°, with 3° increment at a defocus value ranging from -5 to -8 µm. The dataset were collected at a nominal magnification of 19,500 × (an effective pixel size of 4.51 Å) in counting mode, with an accumulated dose of ∼130 e^-^/Å^2^ per tilt series.

The motion correction of tilt series was performed by *alignframes* and tomograms were reconstructed manually at a binning factor of 4 using IMOD package^46^. The contrast of some cryo-tomograms was further improved by *isonet*^47^.

### Sub-tomogram averaging

The extended and contracted AlgoCIS particles on the bacterial surface were manually picked from tomograms using the dipole model in *Dynamo*^48^. Sub-tomograms were cropped from the CTF-corrected tomograms (phase flipped) at a binning factor of 4 using *dtcrop* and their azimuth orientations were randomized using *dynamo_table_randomize_azimuth* before alignment. The cropped sub-tomograms from the extended and contracted AlgoCIS (the extended AlgoCIS: 304 particles from 65 tomograms; the contracted AlgoCIS: 31 particles from 15 tomograms) were first aligned in rough angular search steps imposing 6-fold symmetry. The particles were then split into half-dataset based on the odd-and-even order using *dteo*. Each half-dataset was further aligned against the same reference in fine angular search steps, where different masks were applied individually. The resolution of the final sub-tomogram average was estimated from the averages of half-dataset based on the Fourier shell correlation (FSC)^49^ using *relion_postprocess*^50^. The resolution of the sub-tomogram average of the extended AlgoCIS on the bacterial surface was estimated to ∼60 Å, while the contracted AlgoCIS on the bacterial surface was ∼60 Å. The maps of the free extended AlgoCIS (EMD-11749) and the one on the bacterial surface were lowpass-filtered to 60 Å and then aligned against each other. Both maps were then used to generated the different map by *diffmap* (https://grigoriefflab.umassmed.edu/diffmap).

### CryoEM data collection

The imaging parameters of all cryoEM dataset are summarized in Extended Data Table 1. CryoEM dataset of the contracted AlgoCIS upon GdCl or low pH treatment were collected on a Titan Krios operating at 300 kV and equipped with Quantum LS filter and K3 direct electron detector (Gatan), at a nominal magnification of 81,000 × (an effective pixel size of 1.10 Å) with 1.5 sec total exposure time at a defocus ranging from -1.0 to -3.0 µm. Dataset were collected as movie stacks with different software [the contracted AlgoCIS treated with GdCl: super-resolution mode using SerialEM; the contracted AlgoCIS treated with low pH: counting mode using EPU (Thermo Fisher)]. Each stack contains 50 frames and the accumulated dose was ∼60 e^-^/Å^2^.

The cryoEM dataset of NC-AlgoCIS treated with low pH was collected on a Titan Krios operating at 300 kV and equipped with Biocontinuum filter and K3 direct electron detector, at a nominal magnification of 33,000 × (an effective pixel size of 2.678 Å) with 5.0 sec total exposure time at a defocus ranging from -1.0 to -3.0 µm using SerialEM. The dataset was collected as movie stacks containing 25 frames and the accumulated dose was ∼60 e^-^/Å^2^.

The frames of movie stacks were motion-corrected and were dose-weighted using MotionCor2^51^, where the dataset of the contracted AlgoCIS treated with GdCl was down-sampled at a binning factor of 2. The generated micrographs (the contracted AlgoCIS treated with GdCl: 11,644; the contracted AlgoCIS treated with low pH: 10,748; NC-AlgoCIS treated with low pH: 1,419) were used for the CTF parameter estimation by Gctf and also the downstream image processing.

### CryoEM image processing

All cryoEM image processing were performed using Relion-3.1^52^ . To determine the cryoEM structure of baseplate in the contracted AlgoCIS treated with GdCl, some particles were first manually picked and used for the training in *cryolo*. The trained model was applied for picking and the predicted particles (561,354 particles from 11,644 micrographs) were extracted at a binning factor of 4 and then performed two rounds of 2D classification. The particles selected from good classes were applied to 3D auto-refinement and subsequent 2D classification without sampling. The good particles were then re-extracted at a binning factor of 2, and were performed one round of 3D auto-refinement and then 3D focused classification. The particles from two 3D classes (Class II and V) were re-extracted at a binning factor of 1, and were applied to second round of 3D local refinement and focused classification. The particles selected from the reasonable 3D classes (Class I and III) were used for the 3D local refinement and also CTF refinement. A total of 30,251 particles were used to determine an overall baseplate structure in the contracted AlgoCIS treated with GdCl at a resolution of 3.7 Å assuming 6-fold symmetry (Extended Data Fig. 2a-b).

Structures of different parts of baseplate were further focused refined based on the determined orientations. For the proximal end of sheath, a 3D local refinement was performed with a mask covering the proximal end of sheath and the top part of baseplate, resulting in a 3.4 Å structure applied 6-fold symmetry (Extended Data Fig. 2a-b). For the peripheral wedge connected with the tail-fiber, all symmetry equivalent orientations of particles were generated by symmetry expansion from C6 to C1, and were then used for 3D focused classification with a mask on the tail-fiber at a binning factor of 2. The particles from one class (Class V) were selected and applied to one round of 3D local refinement and focused classification. The particles from the reasonable classes (Class II and III) were pooled and performed 3D local refinement at a binning factor of 1. A total of 72,307 particles were used to determine a 3.9 Å structure of the peripheral baseplate wedge connected with a tail-fiber assuming C1 symmetry (Extended Data Fig. 2a-b).

To determine the cap module in the contracted AlgoCIS, the center of particles was shifted from the baseplate to the cap along Z axis (450 pixels) based on the determined particle orientation. One round of 2D classification without angular sampling was performed to remove the missed aligned particles at a binning factor of 1, and the good particles were applied to one round of 3D local refinement and focused classification. The particles from good 3D classes (Class II and III) were re-extracted with a smaller boxsize and performed a 3D local refinement. The map quality was further improved after CTF refinement. A total of 12,926 particles were used to determine the structure of the cap module in the contracted AlgoCIS at a resolution of 3.2 Å assuming 6-fold symmetry (Extended Data Fig. 2a-b).

To determine the whole contracted AlgoCIS, the center of particles was further shifted from the cap module to the center of particle along Z axis (220 pixels) based on the determined particle orientation. Due to the huge size of complex, the particles were only extracted at a binning factor of 2 (boxsize is 500 pixel) and performed a 3D local refinement. A final structure of the contracted AlgoCIS was determined at a resolution of 4.4 Å (Extended Data Fig. 2a), which reached the Nyquist frequency at a binning factor of 2. We did not further pursue the data processing at a binning factor of 1, because current map quality was sufficient enough to build up the overall model and avoided the potentially exhaustive computational resource.

To determine the cryoEM structure of baseplate in the contracted AlgoCIS treated with low pH, a similar image processing strategy was performed as above. The particles were picked using *cryolo* and then applied to two rounds of 2D classification at a binning factor of 4. After removing bad particles, one round of 3D auto-refinement and local classification were performed, and the particles from reasonable classes (Class I and V) were used for second round of 3D auto-refinement and focused classification at a binning factor of 2. The particles from the selected classes (Class III and IV) were used for 3D auto-refinement and also CTF refinement. A total of 8,771 particles were used to determine a 3.7 Å structure of baseplate in the contracted AlgoCIS treated with low pH assuming 6-fold symmetry (Extended Data Fig. 3b-c).

For the image processing of the NC-AlgoCIS treated with low pH, some particles were manually picked and then were used to 2D classification to generate initial 2D classes for the reference-based auto-picking in Relion. 189,363 particles were predicted after auto-picking and were performed one round of 2D classification at a binning factor of 2. After removing bad particles, one round of 3D auto-refinement was performed. The particles were then re-exacted and the center of particles was shifted to the baseplate ring along Z axis based on the determined orientation. One round of 3D local refinement was performed, showing that two different conformations of baseplate (Class 01: a closed cage; Class 02: a partially opened cage). The particles from the classes of the close cage were performed one round of 3D auto-refinement and downstream 2D classification without angular sampling. The good particles (40,554 particles) were then applied to CTF refinement, resulting a 5.4 Å structure of baseplate in the extended state assuming 6-fold symmetry. To determine the structure of the baseplate with partially open cage, the particles from the selected class were performed 3D auto-refinement and CTF refinement. A total of 15,137 particles were used to determine a resolution of 6.6 Å structure of the baseplate with partially open cage using 6-fold symmetry (Extended Data Fig. 8b/d).

### Magnification calibration

Since the pixel value in the dataset of NC-AlgoCIS treated with low pH was not calibrated precisely, the magnification calibration was performed as previously report^53^ using the extended baseplate structure (PDB-7AEB) as a reference. Briefly, a range of pixel values were applied to the reconstructed map of the extended NC-AlgoCIS using *e2proc3d.py*^54^. Each map with scaled pixel value was converted to hkl file using CCP4 package *Sfall*^55^. A rigid-body search of the extended baseplate structure was performed in reciprocal space using *MolRep*^56^ and Fourier space correlation coefficients (CC) of the best search results were plotted (Extended Data Fig. 8c). The pixel size with maximum CC value was regarded as the calibrated pixel size, which was found at the value of 2.64 Å/pixel.

### Structural modeling

To build the model of the contracted AlgoCIS treated with GdCl, the reconstructed map of different parts in the contracted AlgoCIS were good enough for modeling (Extended Data Fig. 2d). Previous structure of the extended AlgoCIS (PDB-7AEB, 7ADZ, 7AEK) and the Alphafold predicted structures (Alg12 and Alg19) were used as initial references after docking into related maps. The docked structures were manually refined in Coot^57^, and then performed real space refinement with iterative refinements of *phenix.real_space_refine*^58^ and RosettaCM^59^. Structures after each iteration of refinement were manually checked in Coot. The structural statistics of the final models were evaluated using *phenix.molprobity*^58^ (Extended Data Table 1) and the *model vs. map* FSCs were calculated using *phenix.mtrifage*^58^ (Extended Data Fig. 2c).

### Negative-stain image data collection and processing

Negative-stain dataset of different AlgoCIS mutants (NC-AlgoCIS and NC-AlgoCIS-AlgoCIS-Alg11^DPM^) treated with low pH were collected on a Tecnai F20 (Thermo Fisher) operating at 200 kV and equipped with CCD with an effective pixel size of 4.22 Å. The dataset was collected using EPU with 1.0 sec total exposure time at a defocus ranging from -1.0 to - 3.0 µm. The collected micrographs (NC-AlgoCIS treated with low pH: 2,426 micrographs; NC-AlgoCIS-Alg19^DPM^: 1,550 micrographs) were used for CTF parameter estimation via Gctf and the downstream image processing.

The overall image processing was similar to the above cryoEM part. The particles picked by *cryolo* were performed two rounds of 2D classification to remove bad particles. The good particles were applied to 3D auto-refinement and then local classification. The particles from different contraction intermediates were selected and performed one round of 3D auto-refinement to determine the reconstructed maps of different contraction intermediates (state-II: 23,693 particles at a resolution of ∼14 Å; state-III: 18,637 particles at a resolution of ∼14 Å; state-IV: 6,964 particles at a resolution of ∼18 Å). The structures of different contraction intermediates were generated by docking different parts of baseplate in the extended or contracted state into the reconstructed maps.

### Structural analysis of different states in the firing of AlgoCIS

To analyze structural re-arrangements of AlgoCIS upon contraction, we superimposed the models of AlgoCIS in the extended and contracted states in two steps, comprising rotational and translational alignments: 1) in rotational alignment we superimposed two models by aligning the inner tube and the N-terminal domain of the cap protein, because both structural components were present and had the same conformations in both states. 2) in translational alignment we aligned two models along Z axis based on the positions of structural components in the baseplate, including sheath initiator protein and baseplate ring.

To analyze the conformation of the baseplate ring in different contraction intermediates, the superimposed baseplate structures from both extended and contracted states were docked into the related reconstructed maps. The baseplate ring part (residue 801-1024 of Alg11; residue 90-376, 495-662, 717-993 of Alg12) in both states were used to generate simulated maps at a resolution of 20 Å, which were used to calculate the correlation coefficients against the reconstructed maps in ChimeraX (Extended Data Fig. 9b).

## Acknowledgements

We thank ScopeM for instrument access at ETH Zürich. We thank the Functional and Genomic Center Zürich (FGCZ) for mass spectrometry support. Pilhofer Lab members are acknowledged for discussions. We thank Florian Wollweber for constructive suggestions in writing and discussion. M.P. was supported by the Swiss National Science Foundation (31003A_179255 and 310030_212592), the European Research Council (679209 and 101000232), and the NOMIS foundation.

## Contribution

J.X. and C.E. conceived the project; C.E. and E.T. generated all mutant strains; C.E. and J.X. purified samples; J.X. prepared cryoEM/ET samples, collected cryoET data, analyzed the reconstructed tomograms and performed sub-tomogram averaging; J.X. collected and processed cryoEM data, built and refined structural models, and performed structural analysis; J.X., C.E. and M.P. wrote the manuscript; all authors commented on the manuscript.

## Declaration of interest

C.E. and M.P. have two provisional patents pending related to CIS in the United States (application no. 62/768,240 and 62/844,988). The other authors declare that no competing interests exist.

## Data availability

The cryoET sub-tomogram averaged maps have been deposited at the Electron Microscopy Data Bank: EMDB-XXXXX (pre-firing AlgoCIS perpendicularly bound to bacterial sufrace) and EMDB-XXXXX (post-firing AlgoCIS perpendicularly bound to bacterial surface). The cryo-tomograms have been deposited at the Electron Microscopy Data Bank: EMDB-XXXXX (cryo-tomogram of pre-/post-firing AlgoCIS perpendicularly bound to bacterial surface). The cryoEM density maps and the related models have been deposited at the Electron Microscopy Data Bank: EMDB-66211 (PDB entry: 9WSZ, the baseplate iris structure in the post-firing state of AlgoCIS); EMDB-66212 (PDB entry: 9WT0, the tail fiber bound to the baseplate wedge in the post-firing state of AlgoCIS); EMDB-66213 (PDB entry: 9WT1, the cap module in the post-firing state of AlgoCIS); EMDB-66214 (the proximal part of sheath-tube module connected to the baseplate in the post-firing state of AlgoCIS); EMDB-XXXXX (the baseplate iris structure in the post-firing state of AlgoCIS upon low-pH treatment); EMDB-XXXXX (the baseplate iris structure in the pre-firing state of non-contractile sheath AlgoCIS mutant upon low-pH treatment); EMDB-XXXXX (the baseplate iris structure with partially open cage of non-contractile sheath AlgoCIS mutant upon low-pH treatment).

---

**Extended Data Table 1.** CryoEM data statistical analysis.

|  | Contracted AlgoCIS treated with GmCl |  |  |  | Contracted AlgoCIS treated with low pH | NC-AlgoCIS treated with low pH |  |
| --- | --- | --- | --- | --- | --- | --- | --- |
|  | Cap (EMDB-66213, PDB-9WT1) | Baseplate iris structure (EMDB-66211, PDB-9WSZ) | Proximal end of sheath (EMDB-66214) | Peripheral wedge and tail-fiber (EMDB-66212, PDB-9WT0) | Baseplate overall (EMDB-XXXXX) | Baseplate in the extended state (EMDB-XXXXX) | Baseplate with partially open cage (EMDB-XXXXX) |
| **Data collection and processing** |  |  |  |  |  |  |  |
| Nominal magnification |  |  | 81,000 |  |  | 33,000 |  |
| Voltage (kV) |  |  | 300 |  |  | 200 |  |
| Electron exposure (e<sup>-</sup>/Å<sup>2</sup>) |  |  |  | 60 (K3 camera) |  |  |  |
| Defocus range (μm) |  |  |  | 1.0-3.0 |  |  |  |
| Pixel size (Å/pixel) |  |  | 1.1 |  |  | 2.64 (calibrated) |  |
| Symmetry imposed | C6 | C6 | C6 | C1 | C6 | C6 | C6 |
| Final particles (No.) | 12,926 | 30,251 | 30,251 | 72,307 | 8,771 | 40,554 | 15,137 |
| Map resolution | 3.2 | 3.7 | 3.4 | 3.9 | 3.7 | 5.4 | 6.6 |
| FSC threshold |  |  |  | 0.143 |  |  |  |
| **Refinement** |  |  |  |  |  |  |  |
| Map sharpening B factor (Å<sup>2</sup>) | -64 | -54 | -82 | -58 | -52 | -98 | 0 |
| Model composition |  |  |  |  |  |  |  |
| Non-hydrogen atoms | 80766 | 92154 |  | 18756 | - | - | - |
| Protein residues | 11088 | 11406 |  | 2329 | - | - | - |
| Chains | 30 | 12 |  | 4 | - | - | - |
| R.M.S deviations |  |  |  |  |  |  |  |
| Bond length (Å) | 0.015 | 0.017 |  | 0.017 | - | - | - |
| Bond angles (°) | 1.403 | 1.554 |  | 1.619 | - | - | - |
| Validation |  |  |  |  |  |  |  |
| MolProbity score | 1.38 | 1.46 |  | 1.31 | - | - | - |
| Clashscore | 5.81 | 5.17 |  | 2.81 |  | - | - |
| Rotamer outlier (%) | 0.08 | 0.06 |  | 0.05 | - | - | - |
| Ramachandran plot |  |  |  |  |  |  |  |
| Favored (%) | 97.73 | 96.92 |  | 96.42 | - | - | - |
| Allowed (%) | 2.05 | 3.08 |  | 3.58 | - | - | - |
| Outlier (%) | 0.22 | 0.00 |  | 0.00 | - | - | - |
| Masked CC | 0.67 | 0.72 | 0.82 | 0.67 | - | - | - |

**Extended Data Table 2.**
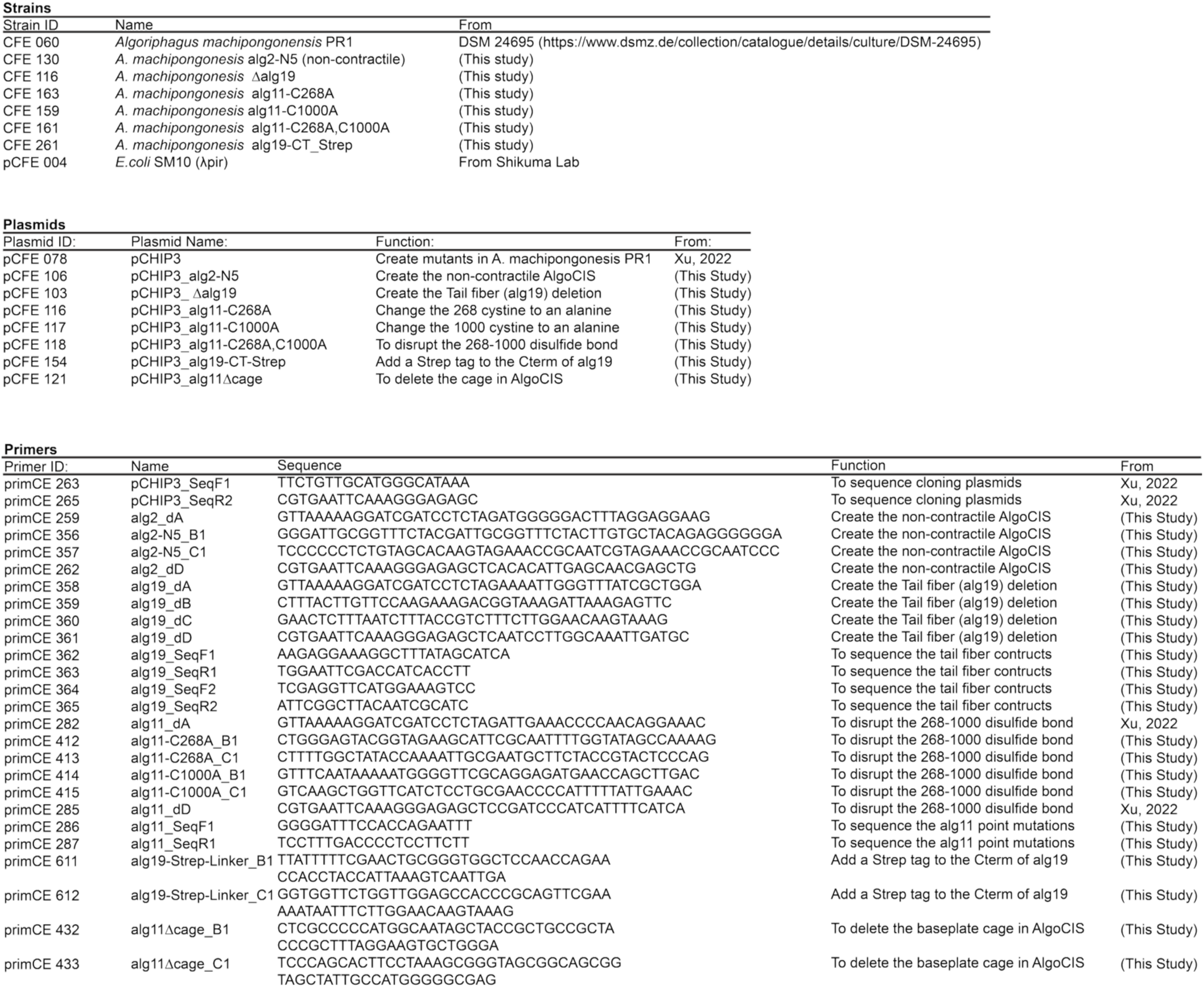
List of strains, plasmids and primers in this study.

## References

1. Costa, T.R.D., Felisberto-Rodrigues, C., Meir, A., Prevost, M.S., Redzej, A., Trokter, M., and Waksman, G. (2015). Secretion systems in Gram-negative bacteria: structural and mechanistic insights. Nat Rev Microbiol 13, 343–359. 10.1038/nrmicro3456.

2. Kooger, R., Szwedziak, P., Böck, D., and Pilhofer, M. (2018). CryoEM of bacterial secretion systems. Curr Opin Struc Biol 52, 64–70. 10.1016/j.sbi.2018.08.007.

3. Shikuma, N.J., Pilhofer, M., Weiss, G.L., Hadfield, M.G., Jensen, G.J., and Newman, D.K. (2014). Marine Tubeworm Metamorphosis Induced by Arrays of Bacterial Phage Tail–Like Structures. Science 343, 529–533. 10.1126/science.1246794.

4. Basler, M., Pilhofer, M., Henderson, G.P., Jensen, G.J., and Mekalanos, J.J. (2012). Type VI secretion requires a dynamic contractile phage tail-like structure. Nature 483, 182–186. 10.1038/nature10846.

5. Lien, Y.-W., Amendola, D., Lee, K.S., Bartlau, N., Xu, J., Furusawa, G., Polz, M.F., Stocker, R., Weiss, G.L., and Pilhofer, M. (2024). Mechanism of bacterial predation via ixotrophy. Sci. (N. York, NY) 386, eadp0614. 10.1126/science.adp0614.

6. Zachs, T., Malit, J.J.L., Xu, J., Schürch, A., Sivabalasarma, S., Nußbaum, P., Albers, S.-V., and Pilhofer, M. (2024). Archaeal type six secretion system mediates contact-dependent antagonism. Sci. Adv. 10, eadp7088. 10.1126/sciadv.adp7088.

7. Rocchi, I., Ericson, C.F., Malter, K.E., Zargar, S., Eisenstein, F., Pilhofer, M., Beyhan, S., and Shikuma, N.J. (2019). A Bacterial Phage Tail-like Structure Kills Eukaryotic Cells by Injecting a Nuclease Effector. Cell Reports 28, 295–301.e4. 10.1016/j.celrep.2019.06.019.

8. Ericson, C.F., Eisenstein, F., Medeiros, J.M., Malter, K.E., Cavalcanti, G.S., Zeller, R.W., Newman, D.K., Pilhofer, M., and Shikuma, N.J. (2019). A contractile injection system stimulates tubeworm metamorphosis by translocating a proteinaceous effector. Elife 8, e46845. 10.7554/elife.46845.

9. Vlisidou, I., Hapeshi, A., Healey, J.R., Smart, K., Yang, G., and Waterfield, N.R. (2019). The Photorhabdus asymbiotica virulence cassettes deliver protein effectors directly into target eukaryotic cells. Elife 8, e46259. 10.7554/elife.46259.

10. Leiman, P.G., Basler, M., Ramagopal, U.A., Bonanno, J.B., Sauder, J.M., Pukatzki, S., Burley, S.K., Almo, S.C., and Mekalanos, J.J. (2009). Type VI secretion apparatus and phage tail-associated protein complexes share a common evolutionary origin. Proc National Acad Sci 106, 4154–4159. 10.1073/pnas.0813360106.

11. Taylor, N.M.I., Raaij, M.J. van, and Leiman, P.G. (2018). Contractile injection systems of bacteriophages and related systems. Mol Microbiol 108, 6–15. 10.1111/mmi.13921.

12. Veesler, D., and Cambillau, C. (2011). A Common Evolutionary Origin for Tailed-Bacteriophage Functional Modules and Bacterial Machineries. Microbiol Mol Biol R 75, 423–433. 10.1128/mmbr.00014-11.

13. Taylor, N.M.I., Prokhorov, N.S., Guerrero-Ferreira, R.C., Shneider, M.M., Browning, C., Goldie, K.N., Stahlberg, H., and Leiman, P.G. (2016). Structure of the T4 baseplate and its function in triggering sheath contraction. Nature 533, 346–352. 10.1038/nature17971.

14. Basler, M., Ho, B.T., and Mekalanos, J.J. (2013). Tit-for-Tat: Type VI Secretion System Counterattack during Bacterial Cell-Cell Interactions. Cell 152, 884–894. 10.1016/j.cell.2013.01.042.

15. Ge, P., Scholl, D., Prokhorov, N.S., Avaylon, J., Shneider, M.M., Browning, C., Buth, S.A., Plattner, M., Chakraborty, U., Ding, K., et al. (2020). Action of a minimal contractile bactericidal nanomachine. Nature, 1–5. 10.1038/s41586-020-2186-z.

16. Sarris, P.F., Ladoukakis, E.D., Panopoulos, N.J., and Scoulica, E.V. (2014). A Phage Tail-Derived Element with Wide Distribution among Both Prokaryotic Domains: A Comparative Genomic and Phylogenetic Study. Genome Biol Evol 6, 1739–1747. 10.1093/gbe/evu136.

17. Chen, L., Song, N., Liu, B., Zhang, N., Alikhan, N.-F., Zhou, Z., Zhou, Y., Zhou, S., Zheng, D., Chen, M., et al. (2019). Genome-wide Identification and Characterization of a Superfamily of Bacterial Extracellular Contractile Injection Systems. Cell Reports 29, 511–521.e2. 10.1016/j.celrep.2019.08.096.

18. Geller, A.M., Pollin, I., Zlotkin, D., Danov, A., Nachmias, N., Andreopoulos, W.B., Shemesh, K., and Levy, A. (2021). The extracellular contractile injection system is enriched in environmental microbes and associates with numerous toxins. Nat Commun 12, 3743. 10.1038/s41467-021-23777-7.

19. Xu, J., Ericson, C.F., Lien, Y.-W., Rutaganira, F.U.N., Eisenstein, F., Feldmüller, M., King, N., and Pilhofer, M. (2022). Identification and structure of an extracellular contractile injection system from the marine bacterium Algoriphagus machipongonensis. Nat Microbiol, 1–14. 10.1038/s41564-022-01059-2.

20. Jiang, F., Li, N., Wang, X., Cheng, J., Huang, Y., Yang, Y., Yang, J., Cai, B., Wang, Y.-P., Jin, Q., et al. (2019). Cryo-EM Structure and Assembly of an Extracellular Contractile Injection System. Cell 177, 370–383.e15. 10.1016/j.cell.2019.02.020.

21. Desfosses, A., Venugopal, H., Joshi, T., Felix, J., Jessop, M., Jeong, H., Hyun, J., Heymann, J.B., Hurst, M.R.H., Gutsche, I., et al. (2019). Atomic structures of an entire contractile injection system in both the extended and contracted states. Nat Microbiol 4, 1885–1894. 10.1038/s41564-019-0530-6.

22. Böck, D., Medeiros, J.M., Tsao, H.-F., Penz, T., Weiss, G.L., Aistleitner, K., Horn, M., and Pilhofer, M. (2017). In situ architecture, function, and evolution of a contractile injection system. Sci New York N Y 357, 713–717. 10.1126/science.aan7904.

23. Casu, B., Sallmen, J.W., Schlimpert, S., and Pilhofer, M. (2023). Cytoplasmic contractile injection systems mediate cell death in Streptomyces. Nat. Microbiol. 8, 711–726. 10.1038/s41564-023-01341-x.

24. Casu, B., Sallmen, J.W., Haas, P.E., Chandra, G., Afanasyev, P., Xu, J., Pilhofer, M., and Schlimpert, S. (2025). Function and firing of the Streptomyces coelicolor contractile injection system requires the membrane protein CisA. bioRxiv, 2024.06.25.600559. 10.1101/2024.06.25.600559.

25. Weiss, G.L., Eisenstein, F., Kieninger, A.-K., Xu, J., Minas, H.A., Gerber, M., Feldmüller, M., Maldener, I., Forchhammer, K., and Pilhofer, M. (2022). Structure of a thylakoid-anchored contractile injection system in multicellular cyanobacteria. Nat Microbiol, 1–11. 10.1038/s41564-021-01055-y.

26. Kreitz, J., Friedrich, M.J., Guru, A., Lash, B., Saito, M., Macrae, R.K., and Zhang, F. (2023). Programmable protein delivery with a bacterial contractile injection system. Nature 616, 357–364. 10.1038/s41586-023-05870-7.

27. Jiang, F., Shen, J., Cheng, J., Wang, X., Yang, J., Li, N., Gao, N., and Jin, Q. (2022). N-terminal signal peptides facilitate the engineering of PVC complex as a potent protein delivery system. Sci Adv 8. 10.1126/sciadv.abm2343.

28. Nachmias, N., Wang, Z., Feng, X., Jiang, F., and Levy, A. (2025). How do bacterial extracellular Contractile Injection Systems bind target cells? A remarkable diversity of receptor binding domains. bioRxiv, 2025.05.13.653841. 10.1101/2025.05.13.653841.

29. Kudryashev, M., Wang, R.Y.-R., Brackmann, M., Scherer, S., Maier, T., Baker, D., DiMaio, F., Stahlberg, H., Egelman, E.H., and Basler, M. (2015). Structure of the Type VI Secretion System Contractile Sheath. Cell 160, 952–962. 10.1016/j.cell.2015.01.037.

30. Ge, P., Scholl, D., Leiman, P.G., Yu, X., Miller, J.F., and Zhou, Z.H. (2015). Atomic structures of a bactericidal contractile nanotube in its pre- and postcontraction states. Nat Struct Mol Biol 22, 377–382. 10.1038/nsmb.2995.

31. Cai, X., He, Y., Yu, I., Imani, A., Scholl, D., Miller, J.F., and Zhou, Z.H. (2024). Atomic structures of a bacteriocin targeting Gram-positive bacteria. Nat. Commun. 15, 7057. 10.1038/s41467-024-51038-w.

32. Marín-Arraiza, L., Roa-Eguiara, A., Pape, T., Sofos, N., Hendriks, I.A., Nielsen, M.L., Steiner-Rebrova, E.M., and Taylor, N.M.I. (2025). Structural characterization of an extracellular contractile injection system from Photorhabdus luminescens in extended and contracted states. bioRxiv, 2025.04.20.649488. 10.1101/2025.04.20.649488.

33. Yap, M.L., Klose, T., Arisaka, F., Speir, J.A., Veesler, D., Fokine, A., and Rossmann, M.G. (2016). Role of bacteriophage T4 baseplate in regulating assembly and infection. Proc National Acad Sci 113, 2654–2659. 10.1073/pnas.1601654113.

34. Abramson, J., Adler, J., Dunger, J., Evans, R., Green, T., Pritzel, A., Ronneberger, O., Willmore, L., Ballard, A.J., Bambrick, J., et al. (2024). Accurate structure prediction of biomolecular interactions with AlphaFold 3. Nature 630, 493–500. 10.1038/s41586-024-07487-w.

35. Park, Y.-J., Lacourse, K.D., Cambillau, C., DiMaio, F., Mougous, J.D., and Veesler, D. (2018). Structure of the type VI secretion system TssK–TssF–TssG baseplate subcomplex revealed by cryo-electron microscopy. Nat Commun 9, 5385. 10.1038/s41467-018-07796-5.

36. Heymann, J.B., Bartho, J.D., Rybakova, D., Venugopal, H.P., Winkler, D.C., Sen, A., Hurst, M.R.H., and Mitra, A.K. (2013). 3-dimensional structure of the toxin-delivery particle antifeeding prophage of Serratia entomophila. Journal of Biological Chemistry 288, jbc.M113.456145-25284. 10.1074/jbc.m113.456145.

37. Nedashkovskaya, O.I., Kim, S.B., Vancanneyt, M., Lysenko, A.M., Shin, D.S., Park, M.S., Lee, K.H., Jung, W.J., Kalinovskaya, N.I., Mikhailov, V.V., et al. (2006). Echinicola pacifica gen. nov., sp. nov., a novel flexibacterium isolated from the sea urchin Strongylocentrotus intermedius. Int. J. Syst. Evol. Microbiol. 56, 953–958. 10.1099/ijs.0.64156-0.

38. Wang, J., Brackmann, M., Castaño-Díez, D., Kudryashev, M., Goldie, K.N., Maier, T., Stahlberg, H., and Basler, M. (2017). Cryo-EM structure of the extended type VI secretion system sheath–tube complex. Nature Microbiology 2, 1507–1512. 10.1038/s41564-017-0020-7.

39. Hu, B., Margolin, W., Molineux, I.J., and Liu, J. (2015). Structural remodeling of bacteriophage T4 and host membranes during infection initiation. Proceedings of the National Academy of Sciences of the United States of America 112, E4919–28. 10.1073/pnas.1501064112.

40. Hu, B., Margolin, W., Molineux, I.J., and Liu, J. (2013). The Bacteriophage T7 Virion Undergoes Extensive Structural Remodeling During Infection. Science 339, 576–579. 10.1126/science.1231887.

41. Leiman, P.G., Chipman, P.R., Kostyuchenko, V.A., Mesyanzhinov, V.V., and Rossmann, M.G. (2004). Three-Dimensional Rearrangement of Proteins in the Tail of Bacteriophage T4 on Infection of Its Host. Cell 118, 419–429. 10.1016/j.cell.2004.07.022.

42. Roderer, D., Bröcker, F., Sitsel, O., Kaplonek, P., Leidreiter, F., Seeberger, P.H., and Raunser, S. (2020). Glycan-dependent cell adhesion mechanism of Tc toxins. Nat. Commun. 11, 2694. 10.1038/s41467-020-16536-7.

43. Tivol, W.F., Briegel, A., and Jensen, G.J. (2008). An Improved Cryogen for Plunge Freezing. Microsc. Microanal. 14, 375–379. 10.1017/s1431927608080781.

44. Wagner, T., Merino, F., Stabrin, M., Moriya, T., Antoni, C., Apelbaum, A., Hagel, P., Sitsel, O., Raisch, T., Prumbaum, D., et al. (2019). SPHIRE-crYOLO is a fast and accurate fully automated particle picker for cryo-EM. Commun. Biol. 2, 218. 10.1038/s42003-019-0437-z.

45. Mastronarde, D.N. (2005). Automated electron microscope tomography using robust prediction of specimen movements. J Struct Biol 152, 36–51. 10.1016/j.jsb.2005.07.007.

46. Kremer, J.R., Mastronarde, D.N., and McIntosh, J.R. (1996). Computer Visualization of Three-Dimensional Image Data Using IMOD. J Struct Biol 116, 71–76. 10.1006/jsbi.1996.0013.

47. Liu, Y.-T., Zhang, H., Wang, H., Tao, C.-L., Bi, G.-Q., and Zhou, Z.H. (2022). Isotropic reconstruction for electron tomography with deep learning. Nat. Commun. 13, 6482. 10.1038/s41467-022-33957-8.

48. Castaño-Díez, D., Kudryashev, M., Arheit, M., and Stahlberg, H. (2012). Dynamo: A flexible, user-friendly development tool for subtomogram averaging of cryo-EM data in high-performance computing environments. J Struct Biol 178, 139–151. 10.1016/j.jsb.2011.12.017.

49. Rosenthal, P.B., and Henderson, R. (2003). Optimal Determination of Particle Orientation, Absolute Hand, and Contrast Loss in Single-particle Electron Cryomicroscopy. J Mol Biol 333, 721–745. 10.1016/j.jmb.2003.07.013.

50. Scheres, S.H.W. (2012). RELION: Implementation of a Bayesian approach to cryo-EM structure determination. J. Struct. Biol. 180, 519–530. 10.1016/j.jsb.2012.09.006.

51. Zheng, S.Q., Palovcak, E., Armache, J.-P., Verba, K.A., Cheng, Y., and Agard, D.A. (2017). MotionCor2: anisotropic correction of beam-induced motion for improved cryo-electron microscopy. Nat Methods 14, 331–332. 10.1038/nmeth.4193.

52. Zivanov, J., Nakane, T., Forsberg, B.O., Kimanius, D., Hagen, W.J., Lindahl, E., and Scheres, S.H. (2018). New tools for automated high-resolution cryo-EM structure determination in RELION-3. Elife 7, e42166. 10.7554/elife.42166.

53. Xu, J., Wang, D., Gui, M., and Xiang, Y. (2019). Structural assembly of the tailed bacteriophage ϕ29. Nat Commun 10, 2366. 10.1038/s41467-019-10272-3.

54. Tang, G., Peng, L., Baldwin, P.R., Mann, D.S., Jiang, W., Rees, I., and Ludtke, S.J. (2007). EMAN2: An extensible image processing suite for electron microscopy. J. Struct. Biol. 157, 38–46. 10.1016/j.jsb.2006.05.009.

55. Winn, M.D., Ballard, C.C., Cowtan, K.D., Dodson, E.J., Emsley, P., Evans, P.R., Keegan, R.M., Krissinel, E.B., Leslie, A.G.W., McCoy, A., et al. (2011). Overview of the CCP4 suite and current developments. Acta Crystallogr. Sect. D 67, 235–242. 10.1107/s0907444910045749.

56. Vagin, A., and Teplyakov, A. (1997). MOLREP: an Automated Program for Molecular Replacement. J. Appl. Crystallogr. 30, 1022–1025. 10.1107/s0021889897006766.

57. Emsley, P., Lohkamp, B., Scott, W.G., and Cowtan, K. (2010). Features and development of Coot. Acta Crystallogr Sect D Biological Crystallogr 66, 486–501. 10.1107/s0907444910007493.

58. Adams, P.D., Afonine, P.V., Bunkóczi, G., Chen, V.B., Davis, I.W., Echols, N., Headd, J.J., Hung, L.-W., Kapral, G.J., Grosse-Kunstleve, R.W., et al. (2010). PHENIX: a comprehensive Python-based system for macromolecular structure solution. Acta Crystallogr Sect D Biological Crystallogr 66, 213–221. 10.1107/s0907444909052925.

59. Song, Y., DiMaio, F., Wang, R.Y.-R., Kim, D., Miles, C., Brunette, T., Thompson, J., and Baker, D. (2013). High-Resolution Comparative Modeling with RosettaCM. Structure 21, 1735–1742. 10.1016/j.str.2013.08.005.

